# Ribosomal protein imbalance launches a C/EBP-based program to preserve tissue integrity

**DOI:** 10.1101/467894

**Authors:** Ludovic Baillon

## Abstract

Genes encoding ribosomal proteins are expressed at rate limiting levels, rendering their biological function highly sensitive to the copy-number variation that results from genomic instability. Cells with a reduced number of ribosomal protein genes (RPGs) are eliminated, when intermingled with wild type cells, via a process known as cell competition. The mechanisms underlying this phenomenon are poorly understood. Here we report the function of a CCAAT-Enhancer-Binding Protein (C/EBP), Xrp1, that is critically required for the elimination of cells with a hemizygous RPG genotype. In such cells, *Xrp1* is transcriptionally upregulated by an autoregulatory loop and is able to trigger cell elimination. Since genomic instability is likely to cause the loss of a haploinsufficient RPG, we propose a molecular model of how RPGs, together with a C/EBP-dependent transcriptional program, could preserve the genomic integrity of tissues.

## Introduction

Multicellular organisms maintain genomic stability via the activation of DNA repair mechanisms to identify and correct damages present in their DNA, cell cycle arrest to prevent the expansion of DNA damaged cells, and finally programmed cell death to eliminate cells irremediably damaged (Ciccia and Elledge, 2010). The P53 transcription factor plays an evolutionary conserved role in the induction of apoptosis following DNA damage, however evidence points towards the existence of alternative routes for the induction of apoptosis in response to DNA damage (McNamee and Brodsky, 2009; Titen and Golic, 2008). It has been proposed that one of these routes relies on the detection of copy number reduction affecting haploinsufficient ribosomal protein genes (*hRPGs*) (McNamee and Brodsky, 2009).

Ribosomes are essential macromolecular machines that catalyze the synthesis of proteins in all cells; they consist of a set of ribosomal proteins (RPs) that surround a catalytic core of ribosomal RNAs (rRNAs). The coordinated function of RPs is well illustrated in *D. melanogaster*. In this model organism, the majority of *RPGs* is haploinsufficient and give rise to the same dominant phenotype referred to as the *Minute* phenotype. This phenotype is characterized by a general developmental delay and improper bristle development (Marygold et al., 2007). Haploinsufficient *RPGs* (*hRPGs*) are widely distributed across the genome, and owing to their dominant adult phenotype these loci have been used to probe the genetic consequences of diverse sources of chromosome damage (Dekanty et al., 2012). This suggests that *hRPGs* may be used to report loss of genetic integrity.

In addition, it has recently been shown that removal of one functional copy of a *hRPGs* activates a number of genes involved in the maintenance of genetic integrity (Kucinski et al. 2017). Furthermore, such prospective loser cells are eliminated when intermingled with wild-type cells, a process that is referred to as “cell competition” (reviewed by Baillon and Basler, 2014). This process occurs irrespectively of the presence of a functional *p53* gene (Kale et al., 2015) and therefore provides a suitable assay to uncover molecular circuitries that can trigger apoptosis in destabilized genomes.

## Results and Discussion

In order to identify genes whose functions are necessary for the elimination of *RPG* heterozygous mutant cells, we performed a mosaic forward genetic screen using ethyl methanesulfonate (EMS) in *D. melanogaster*. We designed a mosaic system that allows direct screening for the persistence of otherwise eliminated loser clones (*RpL19* ^+/−^) through the larval cuticle (Fig. 1A,B). Our F1 genetic mosaic system enabled us to screen for a wide spectrum of suppressors, either dominant mutations (anywhere in the genome) or recessive mutations on the right arm of the third chromosome. In brief, the induction of a single somatic recombination event between two FRTs (FLP recognition targets) generates a *RPG* heterozygous mutant cell that becomes homozygous for the mutagenized right arm of the third chromosome (Fig. 1A). These loser cells/clones are induced at the beginning of larval development (L1). If no suppressive mutation is present, these clones are efficiently eliminated over time such that they are not detectable any more by the end of the third instar larval stage (L3) when the screening is performed (Fig. 1B). This screen should achieve a high degree of specificity since a random mutagenic event is more likely to promote the elimination of a loser cell rather than suppressing it.

**Figure 1:**
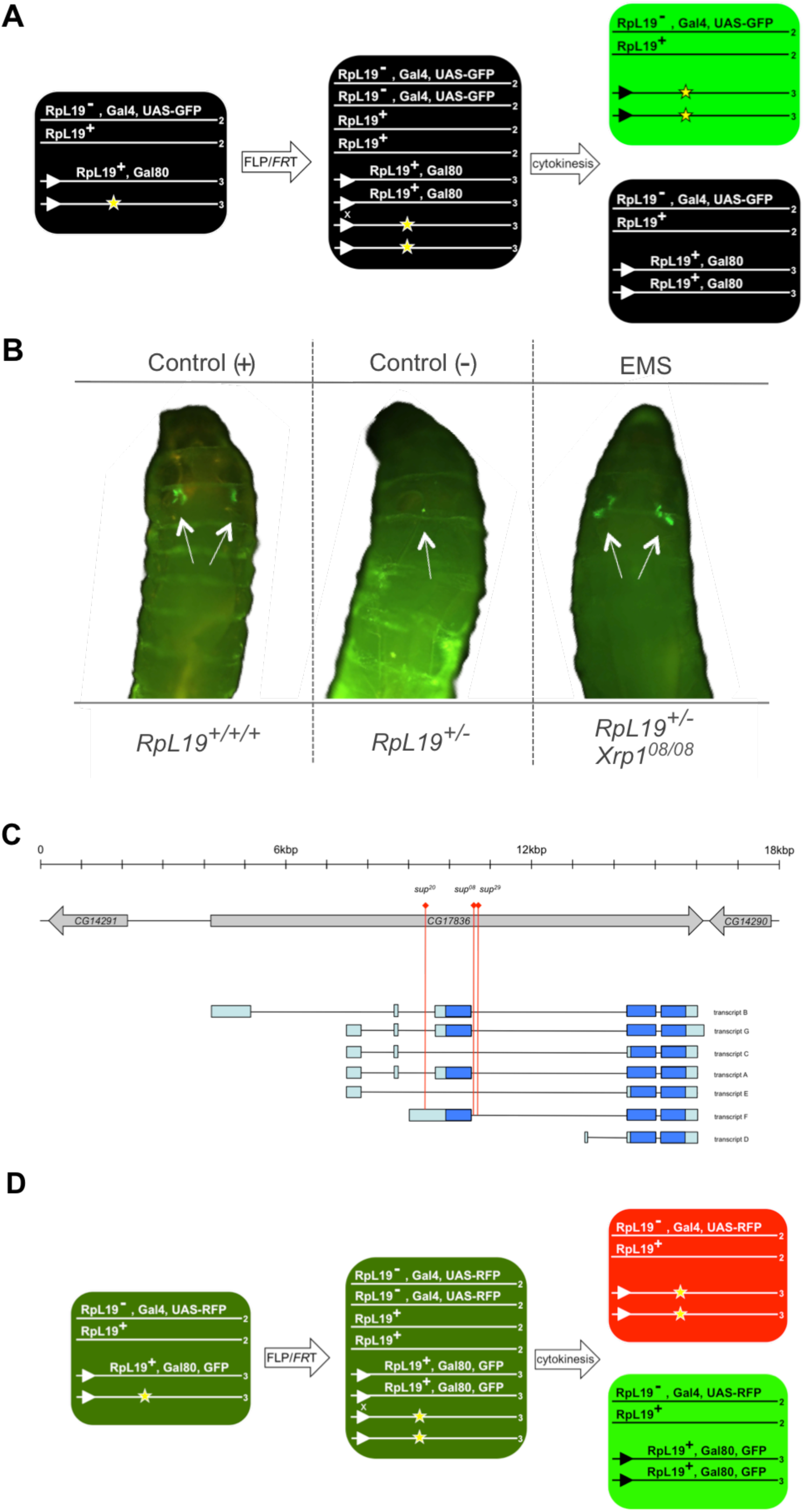
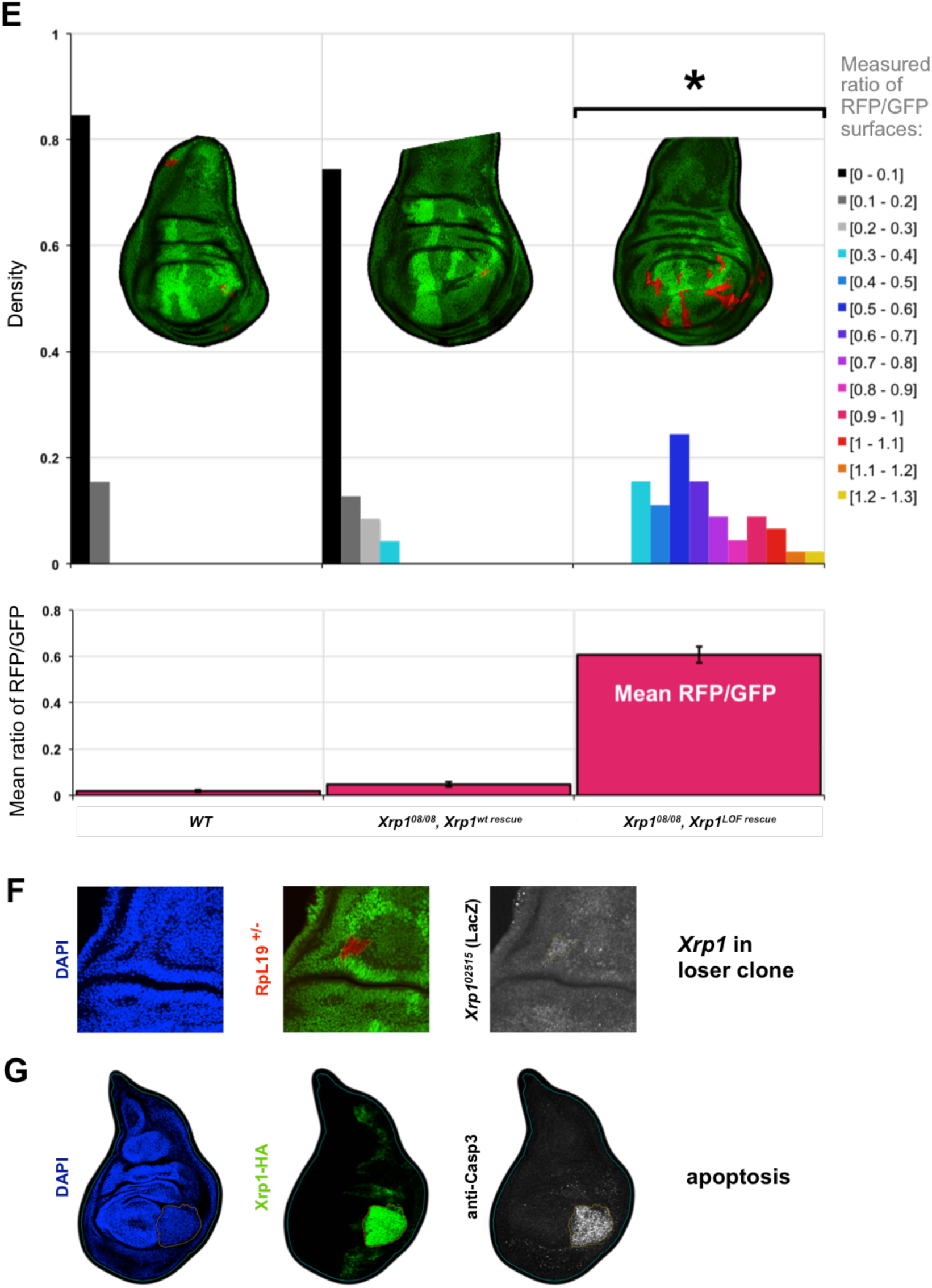
Identification of Xrp1, a gene required for the elimination of *RPG*^+/−^ loser cells. **(A)** Schematic of the genetics used to generate *RpL19*^+/−^ loser clones in a wild-type background using the FLP/*FRT* system. Lines represent chromosomes, numbers at the end of each line indicate the chromosome number, triangles represent the FLP recognition target (FRT) on the right arm of chromosome 3. Site-directed recombination occurs between *FRTs* when the expression of FLP is induced via heat-shock. The yellow asterisk marks the chromosome to be tested for the presence of an EMS induced suppressor. The arrangement depicted here is a variation of the classical MARCM technology that allows us to GFP label cells that are *RpL19*^+/−^ and homozygous for a mutagenized chromosome arm 3R. **(B)** Representative examples of living larvae displaying GFP clones in the pouch of the wing imaginal discs. Gal4 expression is limited to the wing pouch (driven by the *SalmE* enhancer fragment) in order to allow direct screening of living larvae through the cuticle. The genotype of the GFP clones is indicated below the larvae. These genotypes at the left and in the middle have been used to calibrate the screening procedure: (Left) Positive control for clone induction using the *FRT82 RpL19*^+^ chromosome. In this case recombination generates *RpL19*^+/+/+^ cells that are not eliminated. (Middle) Negative control for clone induction using the isogenized *FRT82* chromosome. Recombination produces *RpL19*^+/−^ cells that are efficiently eliminated leaving occasionally one or several small clone fragments in the pouch of the wing disc. (Right) Suppressor *Xrp1^08^* retrieved from the EMS screen gives rise to a consistent GFP signal in both wing pouches. Detailed genotypes for each figure panel are listed in Supplementary Table S3. **(C)** Dmel_r6.08 annotated *Xrp1* transcripts are designated with a single letter from A to G. Transcripts are ordered according to the transcriptional start site and the transcript length. Grey color indicates the gene region, blue the coding regions and light blue the untranslated regions. The red lines indicate the position of the three *Xrp1* alleles retrieved from the EMS screen (*Xrp1^20^*, *Xrp1^08^*, *Xrp1^29^*), none of them is localized in a coding region. (Bottom) The white rectangles represent the genomic fragments used to revert the suppression of cell competition mediated by the allele *Xrp1^08^*. The regions hatched in blue (first and second constitutive exon) within the mutated genomic fragment indicate a shift of the reading frame. **(D)** Schematic of the genetics used to generate *RpL19*^+/−^ loser clones (red) and their respective twin spot (bright green) such that the sizes of these two clones can be compared. Under normal conditions loser clones are eliminated. **(E)** The clone areas of the RFP and of the GFP clones were measured for each individual discs. The upper part of the graph provides the probability density function of the size ratio RFP/GFP of all the discs analyzed. The lower part of the graph indicates in dark magenta the mean RFP/GFP size ratio for each genotype (respectively 0.019, 0.047, 0.607). The sample size of each genotype is respectively 52, 47 and 45 wing imaginal discs. Error bars are SEM (respectively 0.004, 0.011, 0.035). The frameshift in the coding sequence of the mutated *Xrp1* genomic fragment fails to complement *Xrp1^08^* and consequently *RpL19*^+/−^ loser clones persist, see red clones in the representative wing disc. In addition, the absence of loser clone elimination is further evidenced by the normality of the probability density function (D’Agostino & Pearson normality test p=0.0737 > 0.05, the asterisk indicates that normality is accepted). The wild-type *Xrp1* genomic fragment restores the elimination of *RpL19*^+/−^ loser clones. Loser red clones are barely detectable and the probability density function, as it is the case for the WT control, is extremely right skewed (D’Agostino & Pearson normality test p < 0.0001, in both cases normality is rejected). Detailed genotypes for each figure panel are listed in Supplementary Table S3. **(F)** *Xrp1* is up-regulated in *RpL19*^+/−^ loser clones. (Left) DAPI, (Middle) RFP indicates the position of an escaper loser clone, and (Right) anti-β-Gal (lacZ) staining reveals ***Xrp1^02515^*** up-regulation. All panels are maximum intensity projections of Z-stacks. Detailed genotypes for each figure panel are listed in Supplementary Table S3. **(G)** Overexpression of *Xrp1* induces apoptosis. (Left) DAPI, (Middle) GFP indicates the position of the cells misexpressing Xrp1, and (Right) anti-caspase 3 staining reveals the induction of apoptosis. All panels are maximum intensity projections of Z-stacks. Detailed genotypes for each figure panel are listed in Supplementary Table S3.

We screened 20,000 mutagenized genomes for the presence of mutations that would allow loser clones to persist. We retrieved 12 heritable suppressors (Supplementary Table S1) and focused our attention on three of the strongest suppressors that did not display an obvious growth-related phenotype. These suppressors did not belong to a lethal complementation group and the causative mutations were identified using a combination of positional mapping and whole-genome re-sequencing (see Experimental Procedures). Positional mapping placed the three suppressive mutations within a 100 kilobase interval that contains twelve genes. Sanger sequencing of the respective annotated exons did not reveal the presence of any mutations. We therefore complemented our initial mapping with whole-genome re-sequencing and identified three independent mutations in the introns of *CG17836*/*Xrp1* (Fig. 1C, Supplementary Text and Supplementary Fig. S1,2).

A role for *Xrp1* in loser cell elimination has been suggested by genetic association (Lee et al., 2016). Furthermore, *Xrp1* expression is mildly upregulated in prospective loser cells (Kucinski et al. 2017). Despite these observations, the functional relevance of *Xrp1* in cell elimination remains elusive. In order to confirm that these mutations affect the function of *Xrp1* and no other unannotated gene we attempted to rescue these alleles with two different transgenes using a newly designed genetic set-up (see Fig. 1D). The WT genomic fragment restores the elimination of loser cells homozygous for the suppressive mutation *Xrp1^08^* while a mutated genomic fragment with a frame-shift mutation fails to do so (Fig. 1E). Taken together, these results indicate that Xrp1 is necessary for the elimination of cells impaired in ribosome biogenesis.

Using a transcriptional reporter for *Xrp1* (*Xrp1^02515^*, containing a *lacZ* P-element in *CG17836*, Akdemir et al., 2007), we found that *Xrp1* expression is upregulated in *RPG* ^+/−^ cells, indicating that it might play an active role in the elimination of loser cells (Fig. 1F). In order to gain insights into this function we conditionally forced the expression of Xrp1 in the posterior half of the wing discs and observed a massive induction of apoptosis as revealed by anti-cleaved caspase 3 staining (Fig. 1G). We next introduced mutations into the *Xrp1* coding sequence and selected for mutants where Xrp1 activity is impaired (Fig. 2A). One of these mutants, *Xrp1^61^*, contains a frame shift mutation upstream of the *Xrp1* basic region-leucine zipper domain (b-ZIP) (Fig. 2B) that completely abrogates its function (Fig. 2A). *Xrp1^61^* homozygous mutants are viable and display no obvious phenotypes. This null allele was then used to quantitatively assess the suppressive potential of loss of *Xrp1* function on the elimination of *RPG* mutant cells (*RpL19*^+/−^) and to compare it to the potential of other genetic alterations previously implicated in affecting cell competition (Fig. 2C, D, E). We undertook a stringent comparative analysis based on the ratios between the areas of loser (RFP) and winner (GFP) clones (Fig. 2E provides the numerical values of the Fig. 2C). We then analyzed the density distribution across individual samples of the same genotype (Fig. 2E, categorized by 0.1 increments of the ratio between GFP and RFP area). We reasoned that a genuine suppressor of *RPG* mutant cell elimination should not only increase the mean size of *RPG* mutant clones but also restore a normal distribution of *RPG* mutant clones. In the absence of competitive elimination the growth of *RPG* mutant clones is still affected but these clones should follow a size distribution that is comparable to the one followed by WT clones.

**Figure 2:**
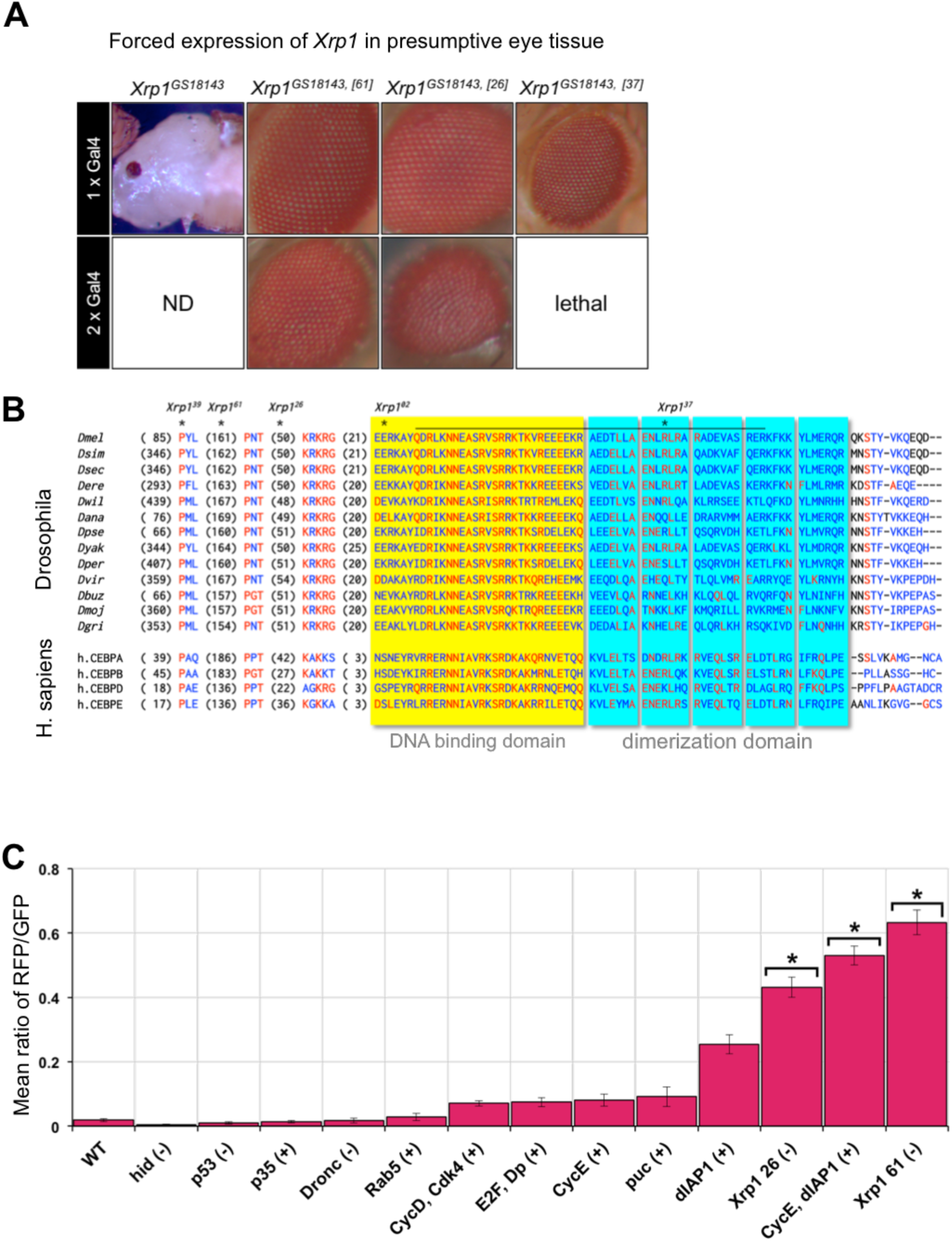
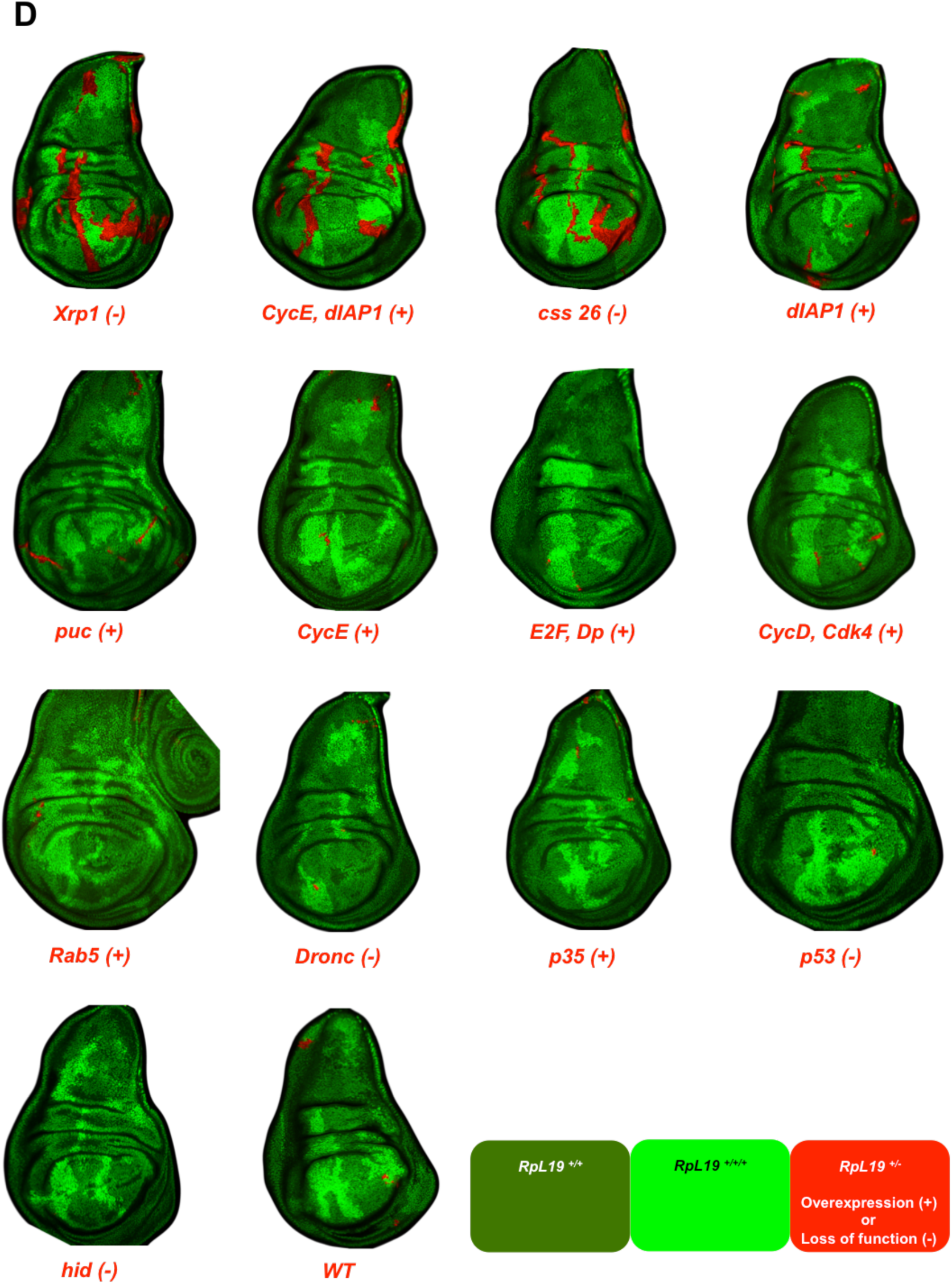
*RPG*^+/−^ loser cell elimination requires multiple inputs. **(A)** Forced expression of *Xrp1* (*Xrp1^GS18143^*) in the presumptive eye tissue is pupal lethal due to the induction of apoptosis in the region where *eyeless* is expressed. EMS induced mutations in the open reading frame of *Xrp1* prevent the induction of apoptosis upon enforced expression of *Xrp1*. *Xrp1^61^* causes a +1 shift of the reading frame after the codon 144 resulting in 60 unrelated amino acids before terminating with two consecutive stop codons. Doubling the dose of the Gal4 driver in *Xrp1^61^* does not lead to detectable *Xrp1* pro-apoptotic activity, thus this allele is considered to be a null allele. Detailed genotypes for each figure panel are listed in Supplementary Table S3. **(B)** Alignment of *Drosophila* Xrp1 protein sequences with the human C/EBPs. Sequences with a minimum of 3 contiguous similar amino acids (AAs) are depicted; red color indicates identity, blue indicates similarity and black means different; the numbers in brackets indicate the length of AAs that do not align. Asterisks indicate the positions of the EMS alleles mapped to the *Xrp1* open reading frame: *Xrp1^39^* (P86S), *Xrp1^61^* (D144>FS), *Xrp1^26^* (Q269•), *Xrp1^02^* (E330K), *Xrp1^37^* (R371•). The line above the *Dmel* Xrp1 sequence delineates the sequence sharing a 40% identity with the human C/EBPs (PSI-BLAST). Highlighted in yellow is the basic region (DNA binding), highlighted in blue is the leucine zipper (hetero/homo-dimerization) of the basic region-leucine zipper domain (b-ZIP). **(C)** Suppressive potential of various genetic alterations as measured by the ratio between the RFP and GFP clone areas. Only *Xrp1* loss of function alleles and the cooverexpression of CycE and dIAP1 are able to substantially rescue loser clone elimination. Asterisks indicate that the distribution of the probability density function does not deviate significantly from normality (see panel E). Detailed genotypes for each figure panel are listed in Supplementary Table S3. **(D)** Representative discs displaying the suppressive potential of various genetic alterations (genotypes are indicated below each discs). Loser clones are labeled with RFP. Detailed genotypes for each figure panel are listed in Supplementary Table S3. **(E)** This panel displays the density function of the size ratio RFP/GFP. Genotypes are ranked according to their suppressive potential. Asterisks indicate the normality (i.e. rescue of clone elimination) as tested by the D’Agostino & Pearson normality test (P values above 0.05 indicate normality). *Xrp1^61^* (p=0.135), *CycE, dIAP1* (p=0.125) and *Xrp1^26^* (p=0.052). For all the other genotypes the probability density functions are extremely right skewed with D’Agostino & Pearson test p values <0.0066. Detailed genotypes for each figure panel are listed in Supplementary Table S3.

Interestingly, unlike loss of *Xrp1*, which fully rescues the elimination of *RpL19*^+/−^ and *RpL14*^+/−^ loser cells (Fig. 2 C, D, E, and Fig. S5), blocking apoptosis by means of overexpression of *dIAP1* or *p35*, or by abrogating *dronc* or *hid* function does not fully suppress *RPG* ^+/−^ cell elimination (Fig. 2C, D, E) suggesting that Xrp1 does more than merely induce apoptosis. Xrp1 may additionally hinder cells to progress through the cell cycle; indeed, *Xrp1* expression has been reported to induce cell cycle arrest in cultured *Drosophila* cells (Akdemir et al., 2007). The co-overexpression of *CycE* (promotes cell-autonomously cell cycle entry (Neufeld et al., 1998) and *dIAP1* (represses cell-autonomously apoptosis (Martin et al., 2009) lead to a suppression of *RPG* ^+/−^ cell elimination comparable to that of loss of *Xrp1* function (Fig. 2C, D, E). This indicates that the combined effects of blocking cell cycle progression and promoting apoptosis are critical for the elimination of *RPG* ^+/−^ mutant cells. Either one or both of these cellular functions could be directly regulated by Xrp1 at the transcriptional level since Xrp1 possesses a sequence-specific DNA binding domain (b-ZIP, Fig. 2B, Fig. S3).

To further explore this notion we set out to identify direct genomic targets of Xrp1 by chromatin immunoprecipitation followed by deep sequencing (ChIP-seq) on wing imaginal discs (Schertel et al., 2015) (see Fig. 3A). ChIP-seq revealed a high number of Xrp1 binding sites spanning a wide range of affinity (see Fig. 3B, C), and establishes Xrp1 as a C/EBP transcription factor (see Fig. 3C, S4). Some of the top targets comprise a number of genes that are involved in cell cycle regulation, apoptosis, and other biological responses previously associated with the elimination of *RPG* ^+/−^ mutant cells (Fig. 3B). Expression analysis at the mRNA (Fig. 3D) or protein level (Fig. 3E and S6) reveals an up-regulation of genes involved in the induction of apoptosis (*hid*, Kale et al., 2015), the regulation of innate immunity (*Dif*, Meyer et al., 2014), and in the establishment of compensatory proliferation (*Upd3*, Kolahgar et al., 2015) as a response to forced Xrp1 expression. Most interestingly, Xrp1 is able to bind to its own promoter and activate its transcription (Fig. 3D, E). The autoregulation of C/EBPs is a feature that is shared among the members of this family of proteins, and it is typically set in motion via mechanisms that are species-specific (Ramji and Foka, 2002). In *Drosophila*, the Xrp1 autoamplification loop could serve as a molecular switch to sustain high transcriptional input on Xrp1 targets to orchestrate the elimination of *RPG* ^+/−^ mutant cells.

**Figure 3:**
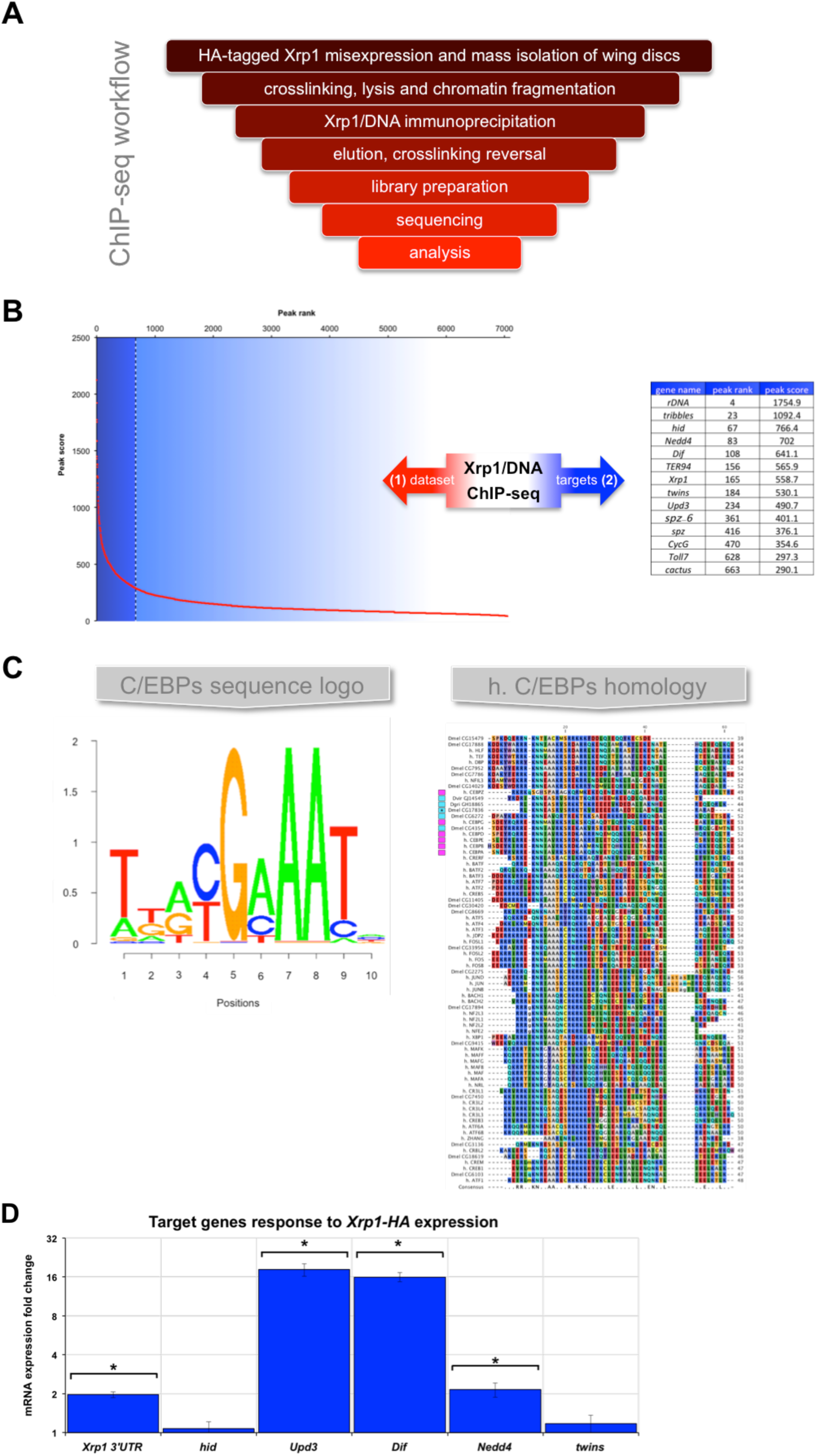
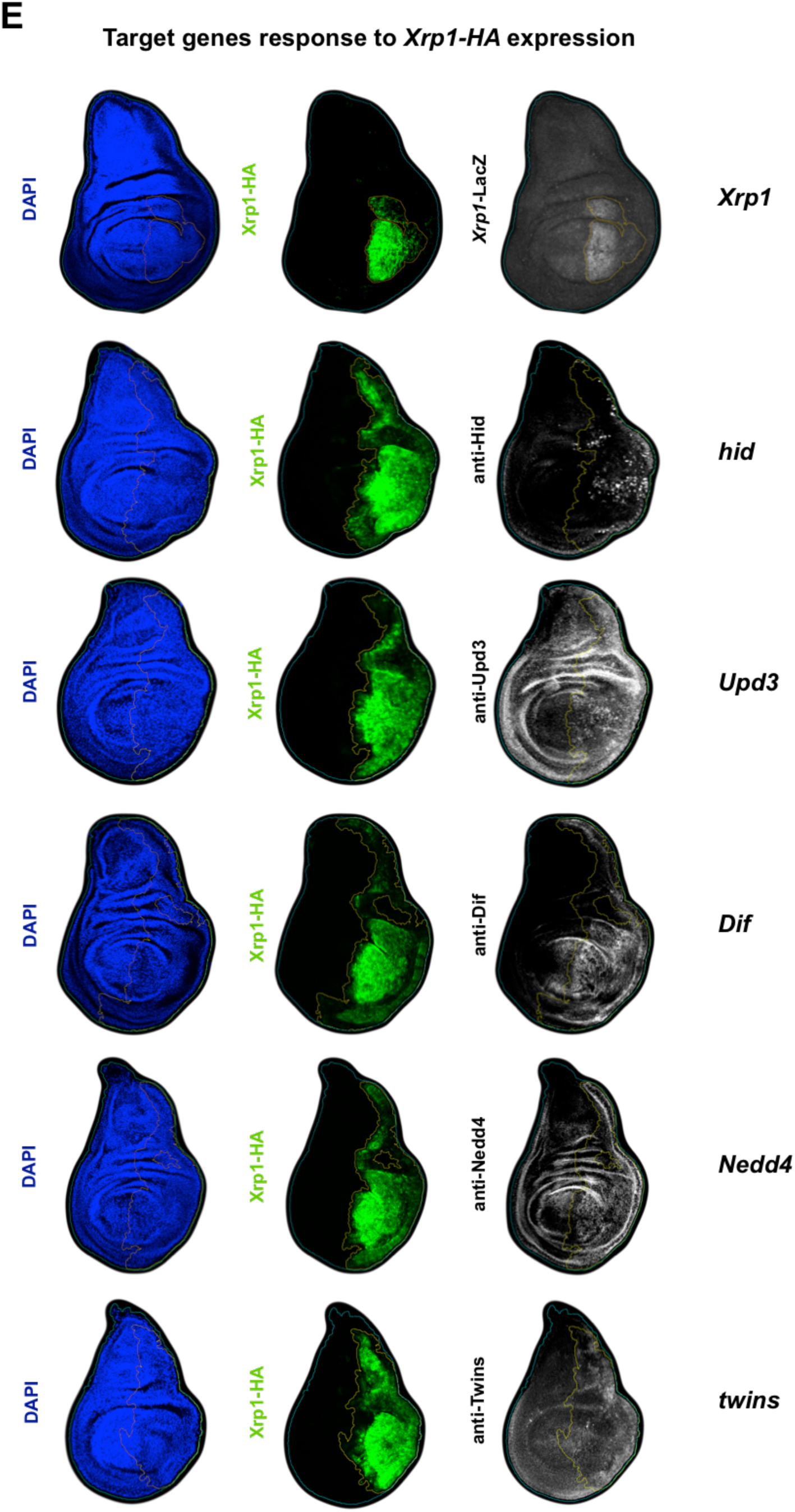
Xrp1 is a CCAAT-Enhancer-Binding Protein (C/EBP) that orchestrates a highly specialized transcriptional program. **(A)** Diagram illustrating the major steps performed in order to identify Xrp1 transcriptional targets via chromatin immunoprecipitation followed by massively parallel DNA sequencing (ChIP-seq). **(B)** (Left) Peak intensity as a function of peak ID. This curve has a typical L shape; a few peaks have high intensities and the vast majority of peaks have comparatively low intensities. The dash line represents the threshold from which targets were no longer considered due to the flattening of the curve. The table on the right lists potential Xrp1 transcriptional targets among Xrp1 genomic target regions having a high peak score. **(C)** (Right) Sequence logo of the most prominent motif bound by Xrp1 (present in 38% of the peaks, p-value ≤ 1e-1009). This motif is related to the b-ZIP binding motif of the C/EBPs protein family. The p-value meets the significance threshold (< 10^−50^). (Left) Seeded/trimmed alignment of all the human and *D. melanogaster* proteins containing the Pfam bZIP_2 motif. Two Xrp1 homologues (*D. virilis* and *D. grimshawi*) were included in the analysis as consistency controls. AAs are colored according to RasMol amino acid color scheme. Colored boxes indicate the human (magenta) and *D. melanogaster* (blue) C/EBPs. The asterisk marks the position of Xrp1. **(D)** Target genes response to Xrp1-HA expression. mRNA expression fold change refers to expression level differences between discs containing clones expressing *Xrp1*-*HA* and discs containing clones expressing a neutral LacZ transcript. Overexpression of Xrp1-HA increases the expression of its genomic counterpart probed via the endogenous 3’ end of the transcripts (*Xrp1 3’UTR*). *Upd3* and *Dif* transcript expression are highly up-regulated by Xrp1 enforced expression. Hid and twins transcript up-regulations are not detectable in this assay. Asterisks indicate statistical significance (t test, p<0.01), errors bars are SEM. **(E)** In this assay, Xrp1-HA expression is temporally enforced in the posterior half of the wing imaginal discs as revealed by the co-expression of GFP. The expression of Xrp1’ targets is probed here by antibody staining. Xrp1 auto-amplification is made visible by the up regulation of *Xrp1*-*lacZ* transcriptional reporter *Xrp1^02515^*. (Left) DAPI, (Middle) GFP and (Right) antibody staining. In this assay, the up-regulation of *hid* is clearly detectable in a subset of cells in the P compartment. The effect of Xrp1-HA expression on *twins* is too subtle to be conclusive. Detailed genotypes for each figure panel are listed in Supplementary Table S3.

C/EBP transcription factors have been shown to prevent cell proliferation and induce apoptosis (Ramji and Foka, 2002). Of particular interest is the retention of C/EBP alpha in the nucleolus via binding to ribosomal DNA (Müller et al., 2010). The nucleolus is a distinct structure in the nucleus of eukaryotic cells that forms around the rDNA. It is the site of ribosome biogenesis and a major stress sensor organelle (James et al., 2014). Cells with only a single functional copy of the *RpL19* gene (*RpL19*^+/−^) have an enlarged nucleolus (Fig. 4A) since RPG insufficiency partially stalls ribosome assembly (Neumueller et al., 2013). Interestingly, Xrp1 binds with a high affinity to many rDNA loci in the genome of *D. melanogaster* (24 peaks mapped, Fig. 3B). The relationship of Xrp1 with the nucleolus and its functional role in the elimination of cells experiencing RPG imbalance points towards a mechanism whereby cells might use the nucleolus as a reservoir of Xrp1 and release it into the nucleus via a nucleolar stress related mechanism. Once in the nucleus, Xrp1 then switches on its own gene via an autoamplification loop and drive cell elimination (a graphical representation of this working hypothesis is presented in Fig. S7).

**Figure 4:**
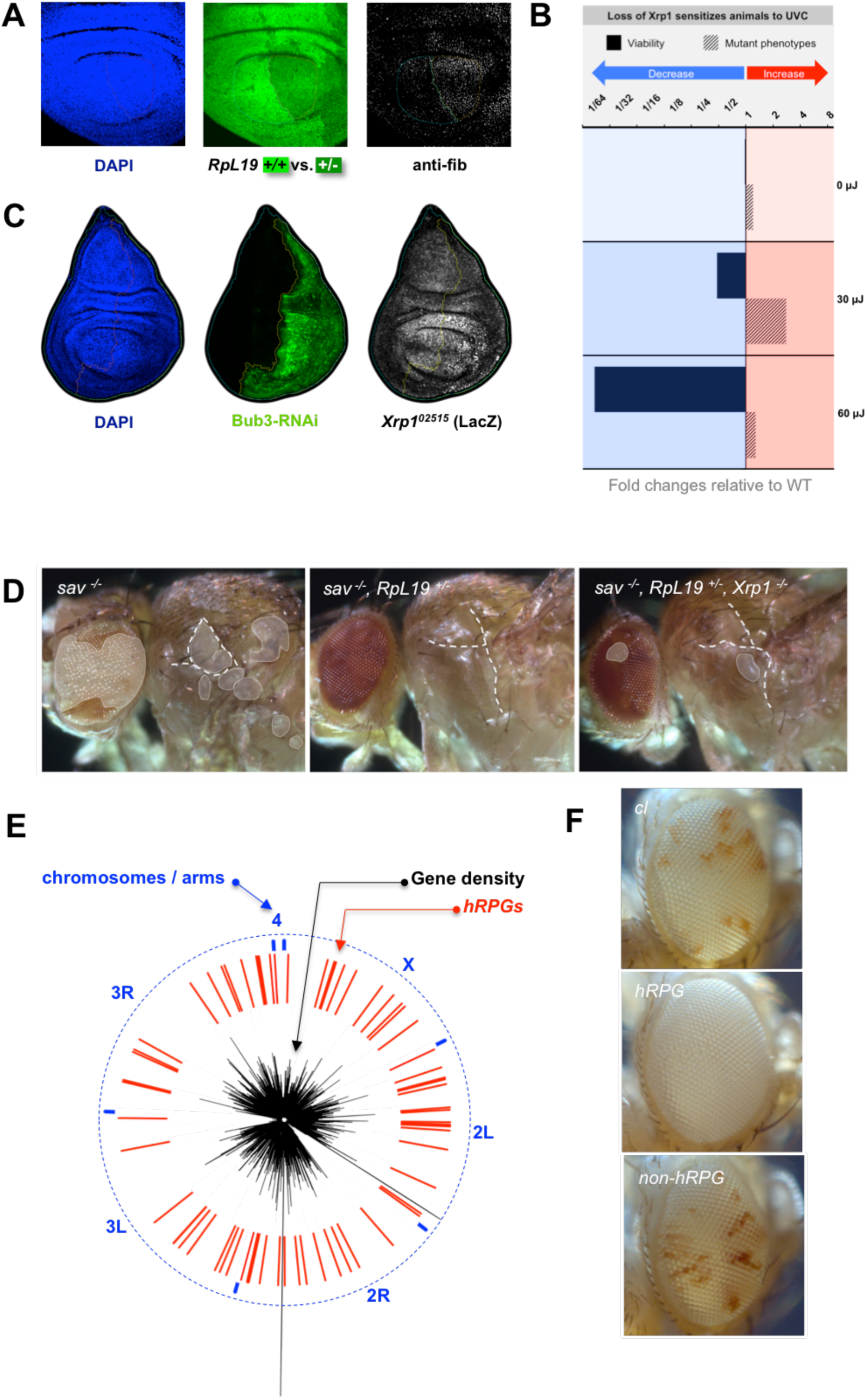
*RPGs* as sensors of genomic integrity. **(A)** Cellular consequences of *RPG* haploinsufficiency. *WT* cells are labeled in bright green and *RpL19*^+/−^ cells are labeled in dark green. *RpL19*^+/−^ cells experience nucleolar stress as revealed by anti-fibrillarin staining. The nucleoli of *RpL19*^+/−^ cells are enlarged due to stalling of ribosome biogenesis. Detailed genotypes for each figure panel are listed in Supplementary Table S3. **(B)** Protective function of Xrp1. Log_2_ ratios of the viability and the mutagenicity of *Xrp1* null animals relative to *WT* exposed to increasing intensity of UV-C (254 nm). As the dose increases, the detrimental effects of UV-C exposure in *Xrp1^61^* animals increases. Black bars represent “viability”, black dashed bars represent “mutant phenotype”, the baseline is set at 1, below the baseline means reduction and above the baseline means increase. Data breakdown is presented in Table S2. **(C)** Xrp1 expression increases upon loss of genomic integrity. Depletion of the spindle assembly checkpoint Bub3 in the posterior half of the wing disc induces chromosome-wide gene dosage imbalances (Clemente-Ruiz et al., 2016) which coincides with an elevated expression of *Xrp1* as revealed by the *lacZ* transcriptional reporter (*Xrp1^02515^*). Elevated expression is detectable in few cells of the posterior compartment. Cells depleted of the spindle assembly checkpoint lose genetic integrity in a stochastic manner. Once the imbalance is established these cells undergo apoptosis and are cleared from the epithelium (Clemente-Ruiz et al., 2016). As a result it is only possible to detect a few cells expressing detectable levels of the *Xrp1* transcriptional reporter. Detailed genotypes for each figure panel are listed in Supplementary Table S3. **(D)** Tumor-suppressive potential of *RPG* haploinsufficiency: heterozygosity for RpL19 is sufficient to fully suppress the emergence of *Salvador* mutant tumors in a *Xrp1* dependent manner. Tumors are outlined and shaded in white, dash white lines indicate the position of the thoracic folds and is used as a point of comparison. Left; *sav*^−/−^ mutant tumors are almost always detectable in adult flies (180/182). Middle; *sav*^−/−^, *RpL19*^+/−^ tumors are never detectable in adult flies (0/158). Right; *sav*^−/−^, *RpL19*^+/−^, *Xrp1*^−/−^ tumors are detectable in adult flies, though notably smaller (171/173). Detailed genotypes for each figure panel are listed in Supplementary Table S3. **(E)** hRPGs are widely distributed across the genome in a way that RP’s imbalance is likely to occur as a result of chromosomal rearrangements. Circular representation of the *D. melanogaster* genome. The blue marks indicate the beginning and the end of the chromosome arms (X, 2L, 2R, 3L, 3R and 4). The black line indicates the relative gene density and the red marks indicate the positions of all haploinsufficient *RPGs*. Detailed genotypes for each figure panel are listed in Supplementary Table S3. **(F)** RPG-induced cell competition is not detectable in non-haploinsufficient *RPG*. *white*^−/−^ clones are generated in *white*^+/−^ background using the Flp/FRT recombination system. The eyeless enhancer drives the expression of FLP that targets recombination between two FRT sites at homologous position on the 3R chromosome arms. (*cl*) Homozygous cells for a cell lethal mutation are not recovered and the eyes are mostly composed of *white*^−/−^ ommatidia. (*hRPG*) Homozygous cells for the *Minute* mutation are not recovered due to lethality and the few heterozygous *white*^+/−^ cells that should remain are eliminated by cell competition. (non-*hRPG*) Homozygous cells for the *non*-*Minute* mutation are not recovered due to lethality but few heterozygous *white*^+/−^ cells remain, indicating the requirement of haploinsufficiency for cell competition.

When intermingled with wild-type cells, cells having only one functional copy of a *hRPG* are eliminated in a *Xrp1*-dependent manner. In our experimental system the deletion of one copy of the *RpL19* gene is catalyzed by the Flp/FRT recombination system which leaves no apparent lesion in the chromosomal DNA (Chen and Rice, 2003). Therefore, the trigger for cell elimination does not depend on DNA damage *per se* but lies within the unbalanced physiology of the cell. The protective function of *Xrp1* at the tissue level and the overall benefit of detecting and eliminating cells with RP imbalance are better illustrated under stress conditions. Abrogation of Xrp1 function sensitizes animal to genotoxic stress such as UV-C (Fig. 4B, Table S2) or gamma rays (Akdemir et al., 2007). Furthermore genomic destabilization following the depletion of the spindle assembly checkpoint gene *bub3* elevates the levels of *Xrp1* expression in the cells that are not yet culled from the epithelia (Fig. 4C and S6). Hence this caretaker mechanism has the potential to preserve genomic integrity at the tissue level by eliminating viable cells that lost genomic integrity. The efficacy of this protective mechanism is probably best illustrated in the context of tumorigenesis for which genomic instability is regarded a major driving force (Hanahan and Weinberg, 2011). We therefore exploited the Flp/FRT recombination system to generate *Salvador*^−/−^ “ mutant tumor clones. In this system, the loss of one functional copy of the *RpL19* gene is sufficient to fully suppress the tumor growth of *Salvador*^−/−^ mutant clones (Fig. 4D; *sav*^−/−^ and *sav*^−/−^, *RpL19*^+/−^). The specific growth arrest mediated by *RpL19*^+/−^ is released upon abrogation of *Xrp1* function (Fig. 4D; *sav*^−/−^, *RpL19*^+/−^, *Xrp1*^−/−^), indicating that the protective function of *RPGs* haploinsufficiency can also operate within tumorous cells. Taken together these results implicate *RPG* haploinsufficiency as a potent tumor suppressor mechanism. In addition to the evidence that loss of *Xrp1* function works as a suppressor of cell competition-driven elimination of both *RpL19*^+/−^ and *RpL14*^+/−^ loser cells (Fig. 2 C, D, E, and Fig. S5), the genome wide distribution of haploinsufficient *RPGs* (*hRPG*) across the *Drosophila* genome (Fig. 4E) further supports the notion that haploinsufficiency at these loci can ensure a prompt response to genomic instability in order to prevent the initiation of tumorigenesis. Haploinsufficiency is the cornerstone of this mechanism since cell elimination is not triggered upon the loss of one functional copy of *RpL3*, a *non*-*hRPG* (Fig. 4F).

## Conclusions

Here we have identified a CCAAT/Enhancer Binding Protein (C/EBP), named Xrp1, as an essential component for the elimination of cells with a reduced copy number of ribosomal protein genes when intermingled with wild type cells. We propose that the imbalanced production of ribosomal proteins triggers a C/EBP dependent transcriptional program that orchestrates the elimination of cells at the onset of genetic instability. Key to this mechanism is the observed haploinsufficiency at RPG loci that translate one-to-one a genetic imbalance into a protein imbalance. This resulting physiological readout at the level of ribosome biogenesis triggers a fail-safe mechanism that leads to the elimination of the impaired cell.

Mutagenesis studies in many diploid organisms indicate that the vast majority of genes are haplosufficient (Wilkie, 1994). Diploidy has been proposed to be evolutionarily selected for as a protective mechanism against the deleterious occurrence of somatic mutations (Orr, 1995). In this view, haploinsufficiency may be considered as an evolutionary accident. In *D. melanogaster*, haploinsufficiency is rare and the vast majority of haploinsufficient genes are *RPGs* (Cook et al., 2012). The haploinsufficiency at these loci is often attributed to the high cellular demand in RPs (Marygold et al., 2007). However, this is not an inherent feature of these genes since there are nine *RPGs* that are not associated with a *Minute* phenotype (*RpSA, RpS2, RpS12, RpL3, RpL23, RpL26, RpL29, RpL30, RpL41*) and for which no functional compensation is possible by paralogous genes (Marygold et al., 2007). The benefits emerging from *RPG* haploinsufficiency (Fig. 4F) appear to outweigh the costs (Orr, 1995) of maintaining it, as it provides a simple, yet effective, mechanism to protect the organism from the emergence of potentially deleterious cells. The basic elements of this mechanism (i.e. *hRPGs* (Barna et al., 2008) at spread-out loci (Uechi et al., 2001), C/EBP (Reinke et al., 2013) and cell competition (Clavería et al., 2013) are conserved, yet it remains to be determined if they function in a similar way in spite of the estimated 600 millions years separating vertebrates from invertebrates. Bearing this in mind, it is tempting to speculate about the therapeutic potential of activating this primitive ribosomal stress response in humans (Fig. 5) since this process occurs irrespectively of the presence a functional *p53* gene (Kale et al., 2015) and since human cancer cells are often mutated for *RPGs* (Fig. S8) (Nijhawan et al., 2012).

**Figure 5:**
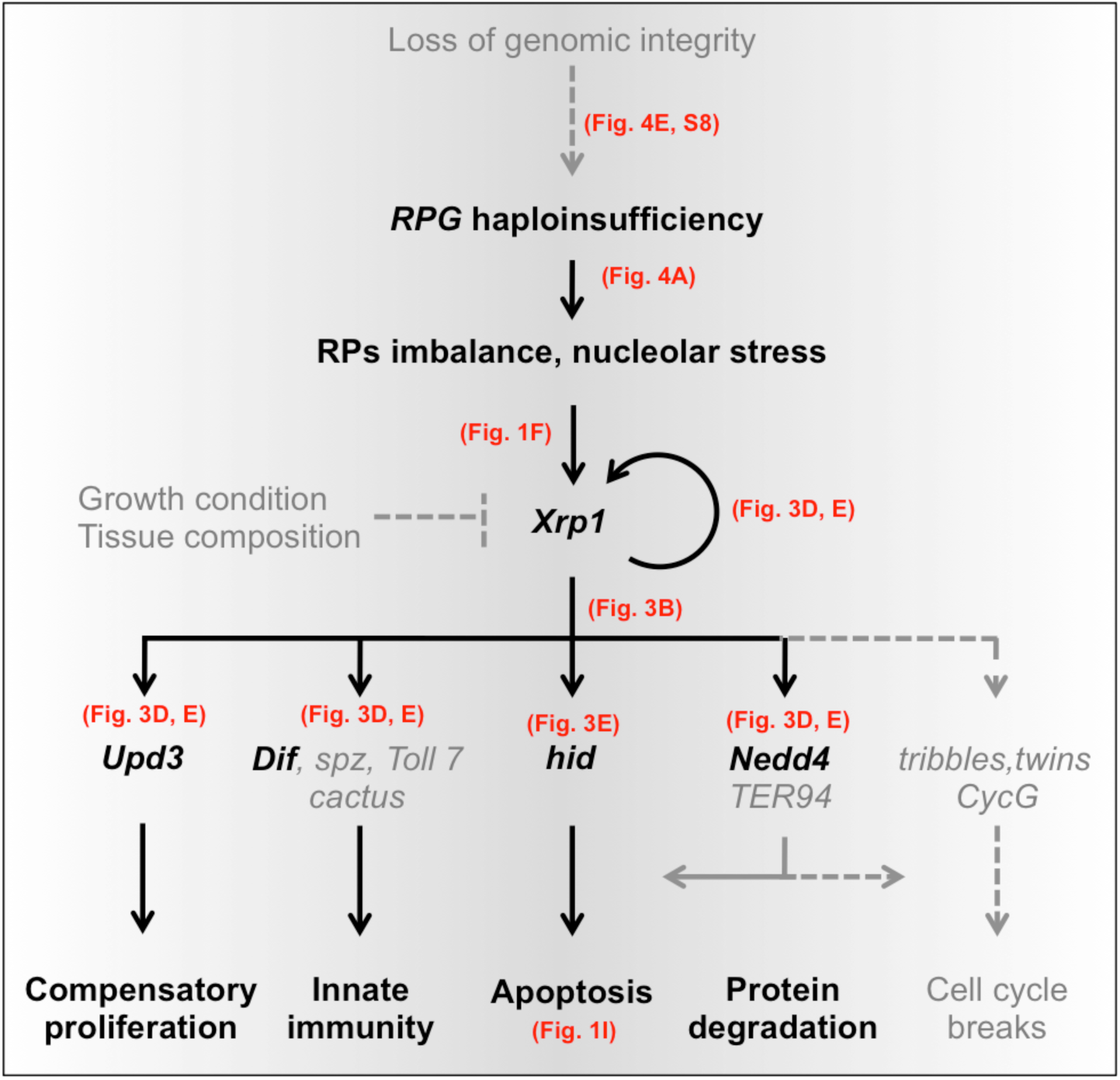
Model for the role Xrp1 may play in maintaining tissue integrity. Model diagram to illustrate the upstream activators and downstream effectors of Xrp1. Arrows indicate activation, bar-headed line indicates repression. Dash lines and grey text indicate parts of the model that could not be thoroughly experimentally confirmed but for which there is strong indirect evidence (i.e. the loss of genomic integrity, the cell non-autonomous layer of regulation upstream or at the level of Xrp1, as well as the cell cycle regulation downstream of Xrp1). Xrp1 activation by ribosomal protein imbalance promotes the elimination and replacement of compromised cells. This active program prevents the emergence of potentially harmful cells that would compromise the integrity of the tissue.

## Experimental Procedures

### *Drosophila* strains and cultures

Flies were grown on a standard cornmeal medium at 25°C unless otherwise specified. The *P{salm*-*GAL4*.*E}2nd* (Denise Nellen, FBrf0211371, 4.8 kbp EcoRI fragment 2L:11459156‥n454345 Dmel_r6.08), the *P{EP}dIAP1* (Chloé Häusermann, line W85 EP insertion at 3L:16046907 Dmel_r6.08) and the *P{en2*.*4*-*GAL4}e16E, P{UAS*-*mCD8::GFP*.*L}LL5, P{tubP*-*GAL80^ts^}* chromosome (Ryohei Yagi) were generated in our laboratory. The *M{UAS*-*Xrp1*.*ORF*.*3xHA*.*GW}ZH*-*86Fb* (F000655) were obtained from FlyORF. The *P{UAS*-*E2F},P{UAS*-*DP}2nd* (#4774), *P{PZ}Xrp1^02515^* (#11569), *P{Act5C*>*y*+>*GAL4*-*w}*, *Df(2R)M60E*, *P{lacW}RpL19^k03704^, P{FRT}82B*, *P{FRT}80B*, *P{tubP*-*GAL80}LL3*, *P{tubP-GAL80}LL9*, *P{Ubi*-*GFP(S65T)nls}3R*, *P{Ubi*-*GFP*.*D}61EF*, the insertion mutants *P{A92}RpS3^Plac92^* (*Minute*) and *P{SUPor*-*P}RpL3^KG05440^* (non-*Minute*) were obtained from the Bloomington *Drosophila* Stock Center. The *P{UAS*-*CycEg}2^nd^* (Lane et al., 1996) and *P{UAS*-*cycD}, P{UAS*-*cdk4}2^nd^* (Meyer et al., 2000) were provided by Christian Lehner. The *P{UAS*-*mCherry*-*CAAX}2^nd^* (Kakihara et al., 2008) was obtained from Shigeo Hayashi. The *P{PZ}hid^05014^*, *P{FRT}80B* and the *Dronc^O1^*, *P{FRT}80B* stocks were provided by Wei Du (Tanaka-Matakatsu et al., 2009). The *p53*^*5A*-*1*-*14*^ corresponds to a 3498 bp deletion within *p53* gene (3R:23,048,029‥23,055,526 Flybase R6.14). The *P{GSV6}Xrp1^GS18143^* (#200976) was obtained from the DGRC Kyoto stock center. Additionally the *P{UAS*-*p35}2^nd^* (Hay et al., 1994), the *P{Rab5}2^nd^* (Entchev et al., 2000) the *P{UAS*-*puckered}2^nd^* (Martín-Blanco et al., 1998), the *sav^4^* (Tapon et al., 2002) were used. *P{Act5C>GAL4*-*w}* was obtained by flipping out the *y*+ FRT cassette of *P{Act5C*>*y*+>*GAL4*-*w}*. The *P{Bub3*-*dsRNA*-*GD9924}2^nd^* was obtained from the VDRC.

### Cloning of transgenes and transgenesis

The *RpL19* 3.08 kbp genomic rescue (2R:24967017‥24970096 Dmel_r6.08) was amplified from a genomic DNA template, sequence confirmed, cloned within the NotI restriction site of the pUAST.attB and inserted into the attP landing site *ZH*-*attP*-*86Fb* (3R tester line) and *ZH*-*attP*-*68E* (3L tester line) (Bischof et al., 2007). The *Xrp1* 15.88 kbp BamHI-BglII genomic rescue (3R:18911505‥18927381 Dmel_r6.08) was digested from CH321-38O16 of the P[acman] BAC Libraries (Venken et al., 2009), sequence confirmed, cloned into the pattB vector (Bischof et al., 2013) and inserted into the *attP* landing site *ZH*-*attP*-*68E*. The *Xrp1* mutated genomic rescue was generated by inserting 5bp (C> GATCCC at 3R:18925226 Dmel_r6.08) at the beginning of the second coding exon in the wild-type genomic fragment, which shifts the frame of all *Xrp1* isoforms. Transgenesis was performed according to standard germ-line transformation procedures.

### RPG loser clone induction and scoring

*RpL19*+/− loser clones *in vivo* screen: *y, w, P{hs*-*FLP}; M{3xP3*-*RFP.attP}ZH*-*36B; P{FRT}82B* mutagenized males were crossed to *y, w, P{UAS*-*mCD8::GFP.L}LL4, P{hs*-*FLP}; P{salm*-*GAL4*.*E}2nd, Df(2R)M60E; P{FRT}82B, P{tubP-GAL80}LL3, M{RpL19 genomic}ZH*-*86Fb*/ *SM5a*-*TM6B* tester virgin females. Parents were allowed to lay eggs for 24 hours and *RpL19*^+/−^ loser clones were heat-shock induced for 30 minutes at 37°C, 24-48 hours after egg deposition. Progeny were screened at the end of the third instar larval stage when larvae stop feeding and move away from the food. No water was added nor was heat-shock applied to force the remaining larvae out of the food as it is routinely done. Special attention was given to the final larval density in the tubes since we noticed that it negatively influences loser clone elimination. In our hands optimal density for cell competition is achieved when the food is neither dry nor soggy; this proper balance is achieved when late third instar larvae climb up to 2/3 of the tube height without reaching the tube’s cotton plug. Consequently only such tubes were screened for the persistence of *RpL19*^+/−^ GFP positive clones through the larval cuticle of living larvae.

*RpL19*^+/−^ loser clones for dissections: males of the appropriated genotype were crossed to the “3R” or “3L” tester virgin females. “3R” tester virgin females: *y, w, P{hs*-*FLP}; P{Act5C>GAL4*-*w}, P{UAS*-*mCherry*-*CAAX}2nd, Df(2R)M60E; P{FRT}82B, P{Ubi*-*GFP(S65T)nls}3R, P{tubP*-*GAL80}LL3, M{RpL19 genomic}ZH*-*86Fb/SM5a*-*TM6B*. “3L” tester virgin females: *y, w, P{hs*-*FLP}; P{Act5C>GAL4*-*w}, P{UAS*-*mCherry*-*CAAX}2nd, Df(2R)M60E; P{Ubi*-*GFP.D}61EF, P{tubP*-*GAL80}LL9, M{RpL19 genomic}ZH*-*68E, P{FRT}80B*/ *SM5a*-*TM6B*. Clones were heat-shock induced as mentioned above.

*RpL14*^+/−^ loser clones for dissections: males of the appropriated genotype were crossed to tester virgin females. (A) *y, w* (B) *y, w, P{hs*-*FLP};; Xrp1^61^* and *y w;; M{UAS*-*RpL14*.*ORF} ZH*-*86Fb* / *TM6B*. Tester virgin females: *y, w, P{UAS*-*mCD8::GFP*.*L}LL4, P{hs*-*FLP};; P{lacW}RpL14^1^, M{salm FRTRpL14 genomic FRT GAL4}ZH*-*86Fb, Xrp1^61^* / *TM6B*. Parents were allowed to lay eggs for 8 hours and loser clones were heat-shock induced for 15 minutes at 37°C, 44-52 hours after egg deposition. Yeast was added to the tubes 24 hours after the heat-shock was applied.

Progeny were screened at the end of the third instar larval stage when larvae stop feeding and move away from the food. No water was added nor was heat-shock applied to force the remaining larvae out of the food.

### Mutagenesis and screen

EMS mutagenesis screens were performed according to standard procedure (Bökel, 2008). *y, w, P{hs*-*FLP}; M{3xP3*-*RFP*.*attP}ZH*-*36B; P{FRT}82B* starter line was first isogenized for the 3R cell competition screen. Isogenized males were fed with a 25 mM EMS, 1% sucrose solution and crossed to tester virgin females. *RpL19*^+/−^ clones were induced in the resulting progeny. A total of 20,000 F1 larvae were screened for the persistence of *RpL19*^+/−^ GFP positive clones at the end of the third instar larval stage. 182 larvae showed persistence of GFP clones clearly above background noise. 125 of them gave rise to fertile adults and were further rescreened. 12 heritable suppressors were doubly balanced. For the *Xrp1* “coding sequence directed mutagenesis” *y, w; +; P{GSV6}Xrp1^GS18143^/*TM3*,Sb* males were fed with a 50 mM EMS, 1% sucrose solution and crossed to tester virgin females *y, w, P{ey-FLP}; P{Act*>*y*+>*GAL4*-*w}; M{3xP3*-*RFP*.*attP}ZH*-*86Fb*. 10,000 F1 genomes were screened and 8 heritable suppressors were retrieved and balanced. A mutation in the *Xrp1* coding region was identified in 5 of them. After the causative mutation was identified the upstream *P{GSV6}Xrp1^GS18143^* was removed using P element transposase and precise excision events were selected (direct sequencing of PCR amplicons) and recombined onto a *P{FRT}82B* chromosome for clonal analysis. *RpL19* knock-out was generated by mobilizing the P element *P{lacW}RpL19^k03704^*, imprecise excisions were selected based on the presence of the characteristic *Minute* bristle phenotype and the absence of the *white*^+^ marker. The *RpL19*^*IE*-*C5*^ 1.09 kbp deletion (2R:24968426‥24969517 Dmel_r6.08) was selected and characterized using direct sequencing of PCR amplicons. This specific excision removes all of *RpL19’s* coding sequence and leaves neighboring genes unaffected.

### Mapping the mutations

We initially mapped cell competition suppressors through meiotic recombinations coupled with DHPLC (Denaturing High-Performance Liquid Chromatography, Eliane Escher) for PCR amplicon analysis. The interval containing the suppressors *Xrp1^08^* and *Xrp1^29^* was narrowed down to a 106.5 Kb interval (3R:18872668‥18979166 Dmel_r6.08). Sanger sequencing of the coding regions in this interval did not reveal the presence of any mutation. We then performed whole-genome sequencing on *Xrp1^08^*, *Xrp1^20^* and *Xrp1^29^* with the Illumina’s Genome analyser IIx (Genomics Platform of the University of Geneva). Mutations were identified by visual inspection of the sequences in this interval: *Xrp1^08^* (T>A 3R:18921364 Dmel_r6.08), *Xrp1^20^*(C>T 3R:18920194 Dmel_r6.08), *Xrp1^29^*(G>A 3R:18921450 Dmel_r6.08). Other suppressors were roughly mapped to the second chromosome or to one of the arms of the third chromosome as indicated in the test complementation table. *Minute* mutants were identified on the basis of their characteristic bristle phenotype and developmental delay. *warts* and *Salvador* mutants were identified on the basis of their clonal overgrown phenotypes and failure to complement independent loss of function alleles (*warts^m72^* and *sav^4^*). Note that for *sup^88^* the suppressive mutation is the *Minute* on the second chromosome and not the mutation in the *salvador* gene. *Xrp1* suppressors isolated from the “coding sequence directed mutagenesis” were identified by direct sequencing of PCR amplicons: *Xrp1^02^* (G>A 3R:18926271 Dmel_r6.08), *Xrp1^26^* (C>T 3R:18926088 Dmel_r6.08), *Xrp1^37^* (C>T 3R:18926394 Dmel_r6.08), *Xrp1^39^* (C>T 3R:18925431 Dmel_r6.08), *Xrp1^61^* (TC>ACA 3R:18925609‥18925610 Dmel_r6.08).

### qRT-PCR

qRT-PCR was performed according to standard protocol. Supplementary Figure S2: RNA was extracted with TRIzol Reagent and genomic DNA was digested with the Ambion DNase kit. RNA was isolated from third instar wing imaginal discs with the exception of the reaction 1-6 where RNA was extracted from third instar larvae. Primer sequences are oriented 5’ to 3’. Primer 1(GCGTAGCAGAAAAGACAAGTGA), 2(CGACACAAGTTCCC CTTAAAC), 3(TCATTGTTTCTTTCTAACGGTCAA), 4(GGTTGCTGTTGTTTG ATTCG), 5(CCTACTGCCACAGTTGAAGAGATAGACG), 6(TTGCTTCTATGT CTTGCAGGTATT), 7(GACCACACCGGAGATTATCAA), 8(GCTGGTACTGGT ACTTGTGGTG).

Figure 3D: RNA was extracted with TRIzol Reagent and genomic DNA was digested with the Invitrogen DNase kit. RNA was isolated from third instar wing imaginal discs. Target gene expression levels were measured in y, w, hsFlp;; Act5C>CD2, y+>GAL4, UAS-GFP / UAS-Xrp1 wing discs and compared to y, w, hsFlp;; Act5C>CD2, y+>GAL4, UAS-GFP / UAS-LacZ control wing discs. GAL4 expressing clones were induced 4 days AED with a 45 minutes heat shock at 37°C. Wing discs were dissected 24 hours after clone induction. Primer sequences are oriented 5’ to 3’. Xrp1 3’UTR Fw (CGTTGAAGAAGTCGAGAAGCA), Xrp1 3’UTR Rev (TAAACACTCCTCGCGCACTA), hid Fw (GTGGAGCGAGAACGACAAA), hid Rev (TTGGCCAAGTGAAGCTCTGT), Upd3 Fw (CCCAGCCAACGATTTTTATG), Upd3 Rev (TGTTACCGCTCCGGCTAC), Dif Fw (GTGGAGCTGAAACTAGTGAGACC), Dif Rev (GGCGATTGTGTTTGGTTAGG), Nedd4 Fw (GACCCTGGTGAATCTGCCTA), Nedd4 Rev (CCGGATAAAGGCGTGGTAG).

### ChIP-seq preparation and analysis

*Drosophila* wing imaginal discs expressing HA tagged *Xrp1* (FlyORF-F000655) (Bischof et al., 2013) were mass isolated/sorted, chromatin was immunoprecipitated and DNA libraries were prepared according to standard protocol (Schertel et al., 2015). Libraries were sequenced on the Illumina HiSeq 2500 v4 (Functional Genomics Center of the University of Zurich). Bowtie 2 (version 2.0.0-beta6) (Langmead and Salzberg, 2012) was used to align the sequencing reads using default parameters. The dm3 *Drosophila* genome annotation was used as reference. The program findPeaks.pl with default parameters was used to identify enriched regions compared to the untreated control sample. The program find MotifsGenome.pl (with the option size =75) was used to identify predominant motifs *de novo*. Only the one motif very significantly enriched (p-value ≪ 1e-50) was considered as advised in the software description. The sequence logo was generated with the PWM-Tools web interface (http://ccg.vital-it.ch/pwmtools/) from the SIB using HOMER’s position frequency matrix output file. Position (A C G T): 1 (0.252 0.046 0.078 0.624), 2 (0.178 0.089 0.344 0.389), 3 (0.468 0.001 0.362 0.169), 4 (0.001 0.560 0.011 0.428), 5 (0.001 0.001 0.997 0.001), 6 (0.590 0.275 0.001 0.134), 7 (0.997 0.001 0.001 0.001), 8 (0.997 0.001 0.001 0.001), 9 (0.082 0.213 0.001 0.704), 10 (0.305 0.287 0.176 0.232).

### Data access

ChIP-seq data and whole-genome resequencing data from this study will be submitted to the NCBI Gene Expression Omnibus data repository.

### Antibodies

Inmunostainings on *Drosophila* wing discs were performed according to standard protocol. The following antibodies were used: rabbit anti-Cleaved-Caspase-3 (Asp175, Cell Signaling), mouse anti-β-Galactosidase (Z3781, Promega), mouse anti-Fibrillarin (38F3; Santa Cruz). The rabbit anti-Dif antibody was obtained from Ylva Engström, the monoclonal mouse anti-Hid antibody was obtained from Hermann Steller, the rabbit anti-Nedd4 antibody was obtained from Shigeo Hayashi, the rat anti-Twins antibody was obtained from Tadashi Uemura and the rabbit anti-Upd3 antibody was obtained from Yu-Chen Tsai. The following secondary antibodies were used: goat anti-mouse Alexa Fluor 647, goat anti-rabbit Alexa Fluor 647 and goat anti-rat Alexa Fluor 647 (Molecular Probes). For the chromatin IP the rabbit anti-HA ChIP grade antibody (ab9110, Abcam) was used. Image processing and clone size measurements were done with FIJI.

### Generation of imaginal wing discs with *RpL19*^+/−^ and *RpL19*^+/+^ compartments

Genotype: *y, w, P{hs*-*FLP}; P{Act5C>GAL4*-*w}, P{UAS*-*mCherry*-*CAAX}2nd, Df(2R)M60E*/*RpL19*^IE-*C5*^ *; P{FRT}82B, P{Ubi*-*GFP(S65T)nls}3R, P{tubP*-*GAL80}LL3, M{RpL19 genomic}ZH*-*86Fb/P{FRT}82B*. Such larvae were heat-shocked 15 min at 37°C during L1. Wing discs were dissected at the end of the third instar larval stage, fixed and stained. Wing discs containing *RpL19*^+/−^ and *RpL19*^+/+^ compartments were imaged.

### Identification of Xrp1 homologs

Heuristic approach. Two iterations of PSI-BLAST (Altschul et al., 1997) were performed using the bZIP domain of Xrp1 as a query. The COBALT constraint-based multiple protein alignment tool provided on the BLAST interface (Papadopoulos and Agarwala, 2007) was used to align all *Drosophila* Xrp1 protein sequences with the human C/EBPs family members identified with the PSI-Blast search. Non-heuristic approach. BZip containing proteins from human and *D. melanogaster* were searched, aligned and trimmed according to the bZIP_2 motif from Pfam (PF07716) using probabilistic hmmer profiles (Eddy, 1998) (hmmer.org). The resulting alignment was visualized with the CLC Main Workbench and then used for phylogenetic reconstruction using the PhyML algorithm (Guindon and Gascuel, 2003) with LG substitution models (Le and Gascuel, 2008), SPR topological rearrangements (Hordijk and Gascuel, 2005) and 100 bootstrap replicates. Phylogenetic tree was then midpoint rooted and displayed with the iTOL online tool (Letunic and Bork, 2007).

### UV-C treatment

L1 larvae (*yw* and *Xrp1^61^*) were exposed to different intensity of UV-C (254 nm) on apple-agar plates and then transferred to standard cornmeal medium. Percentage of eclosed animals was scored as well as the percentage of adults exhibiting obvious phenotypes (bristle number/morphology, eye/notum/wing malformations and melanotic masses).

### RPGs cancer-deletion profiling

The RPG cancer-deletion analyses was performed for each of the 79 human RPGs using the TUMORSCAPE query function based on the GISTIC analysis (Mermel et al., 2010). Deletions for RPGs were significantly detected within the 14 cancer subtypes that regroups more than 80% (2549/3131) of the cancer samples on which the GISTIC analysis is based. This analysis identifies genes that are significantly deleted across the datasets and may therefore underestimate the extend of RPGs deletion due to intra-tumour heterogeneity. Cancer subtypes with GISTIC-based RPG deletions include: Acute lymphoblastic leukemia (391 cancer samples, including 13 cell lines), Breast (243 cancer samples, including 50 cell lines), Colorectal (161 cancer samples, including 33 cell lines), Esophageal squamous (44 cancer samples, including 12 cell lines), Glioma (41 cancer samples, including 13 cell lines), Hepatocellular (121 cancer samples, including 11 cell lines), Lung NSC (733 cancer samples, including 105 cell lines), Lung SC (40 cancer samples, including 23 cell lines), Medulloblastoma (128 cancer samples, including 9 cell lines), Melanoma (111 cancer samples, including 108 cell lines), Myeloproliferative disorder (215 cancer samples, including 0 cell lines), Ovarian (103 cancer samples, including 7 cell lines), Prostate (92 cancer samples, including 9 cell lines), Renal (126 cancer samples, including 27 cell lines).

### *Drosophila RPGs* map / gene density

Gene coordinates for each chromosome arm were retrieved using the cytosearch tool of Flybase. Gene positions were considered as the middle point between the start and the end of each gene. Gene density was calculated for 40 kbp bins and the final map was visualized using the radar chart type of Excel. The percentage of intragenic sequences was calculated as the complement of the total size of the genome minus the sum of the intergenic sequences downloaded from Flybase (Genome, FTP, r6.1).

## Acknowledgments

I would like to thank Konrad Basler for offering me the possibility to join his laboratory and for being such a kind-hearted PhD advisor. I am thankful to Claudia Rockel for carrying out the Chromatin IP, the libraries preparation and for handling the ChIP-seq data, to Jochen Hilchenbach for performing the RT-PCR experiments (quantification of target gene expression), and to Federico Germani for outlining the areas of the clones and performing the test with the RpL14 flip-out cassette. I wish to thank Johannes Bischof for invaluable comments on earlier versions of this manuscript, Eliane Escher (DHPLC genotyping, Sanger sequencing), Nellcia Wang (mass isolation and sorting of wing imaginal discs for the ChIP-seq). I am grateful to Christian von Mering for his guidance on phylogenetic tree construction and to Werner Boll for his help with confocal microscopy. I am grateful to Shigeo Hayashi, Christian Lehner, Wei Du, Marco Milan, Hermann Steller, Ylva Engström, Tadashi Uemura, Shigeo Hayashi, Yu-Chen Tsai for sharing reagents and ideas. I apologize to the researchers whose work could not be cited due to space limitations. This work was supported by the Swiss National Science Foundation and the University of Zurich.

## Supplementary Information

### Xrp1 intronic mutations

The intronic mutations we identified in our screen are all substitutions of single nucleotides. These nucleotides are conserved within the *Drosophila* genus and inspection of the alignment revealed an embedment of these nucleotides in conserved putative motifs (supplementary Figure S1). Of particular interest are the polypyrimidine motifs containing the nucleotide mutated in *Xrp1^20^* and *Xrp1^08^*. These motifs flank the alternative first exon and are potential splice regulators. The CTCTCT motif present near the 5’ splice site of *Xrp1* has been identified as a putative intronic splicing enhancer (ISE) in *Drosophila* (Brooks et al., 2011). This motif is one of the two ISEs that is conserved in vertebrates and is predicted to be a binding site for the polypyrimidine-tract binding protein (PTB) splicing regulator (Brooks et al., 2011). The presence of these motifs prompted us to investigate the consequences of the *Xrp1^08^* allele on exonic junctions. The most prominent effects of this allele are a strong and consistent reduction in the expression of two similar *Xrp1* transcripts, RC and RE (supplementary Figure S2). These transcripts differ only in the composition of their 5’UTRs. They share the same transcriptional start site and contain the same protein-coding sequences that code for the short isoform of this gene (supplementary Figure S1).

**Figure S1.**
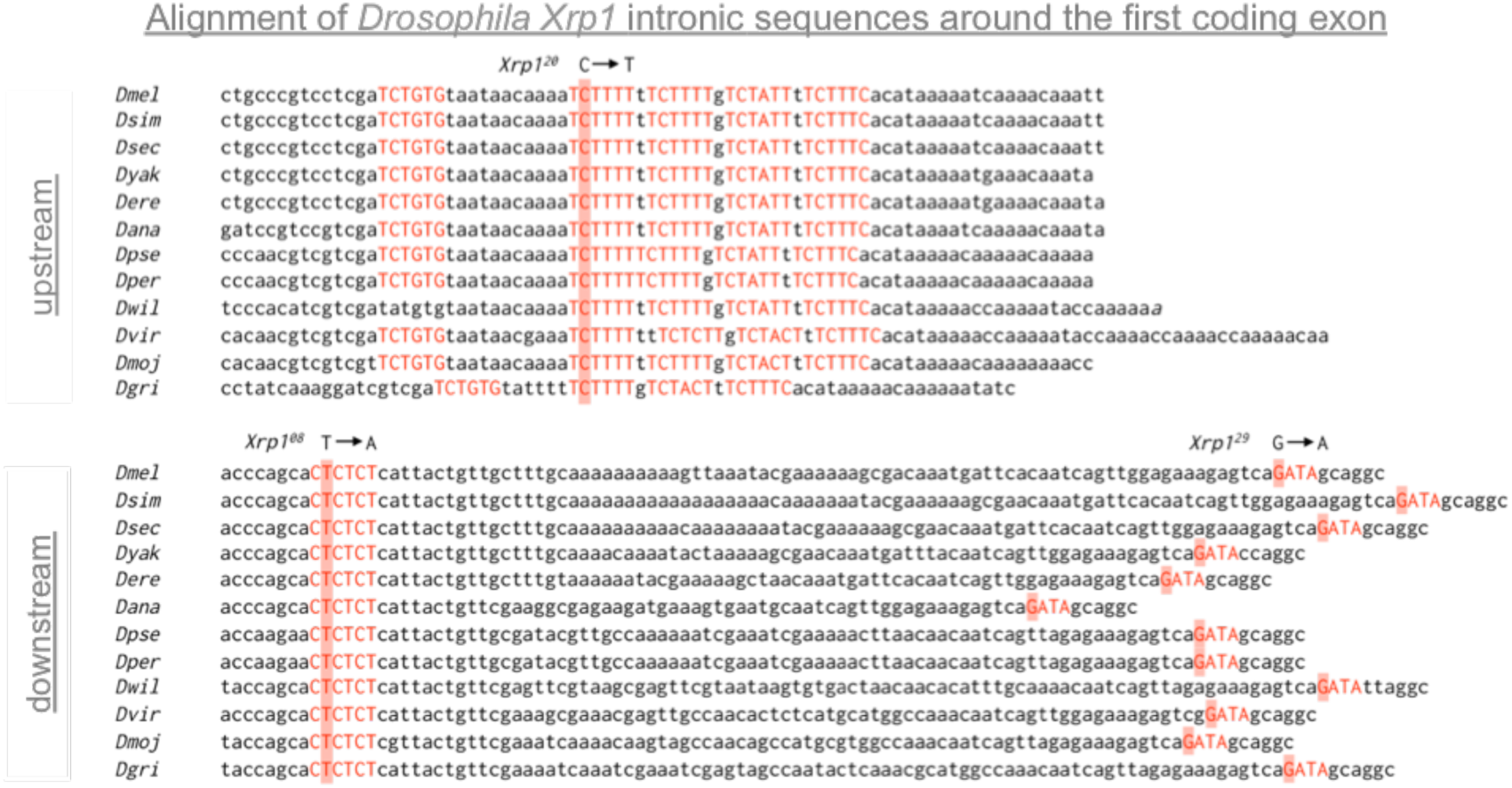
Alignment of Drosophila *Xrp1* sequences. The mutations retrieved from the EMS screen are highlighted in red. All these alleles are single base pair substitutions of conserved nucleotides. The nucleotide substitutions are indicated on top of the *Drosophila melanogaster* sequence (*Dmel*). Conservation extends beyond these nucleotides and seems to affect intronic motifs. These motifs are capitalized and colored in red. The allele *Xrp1^20^* disrupts the repetition of the conserved hexanucleotide pyrimidine motif TCTDTB (D stands for A, G or T and B stands for C, G or T). The allele *Xrp1^08^* disrupts the conserved putative intronic splice enhancer (ISE) CTCTCT and *Xrp1^29^* disrupts a conserved GATA motif

**Figure S2.**
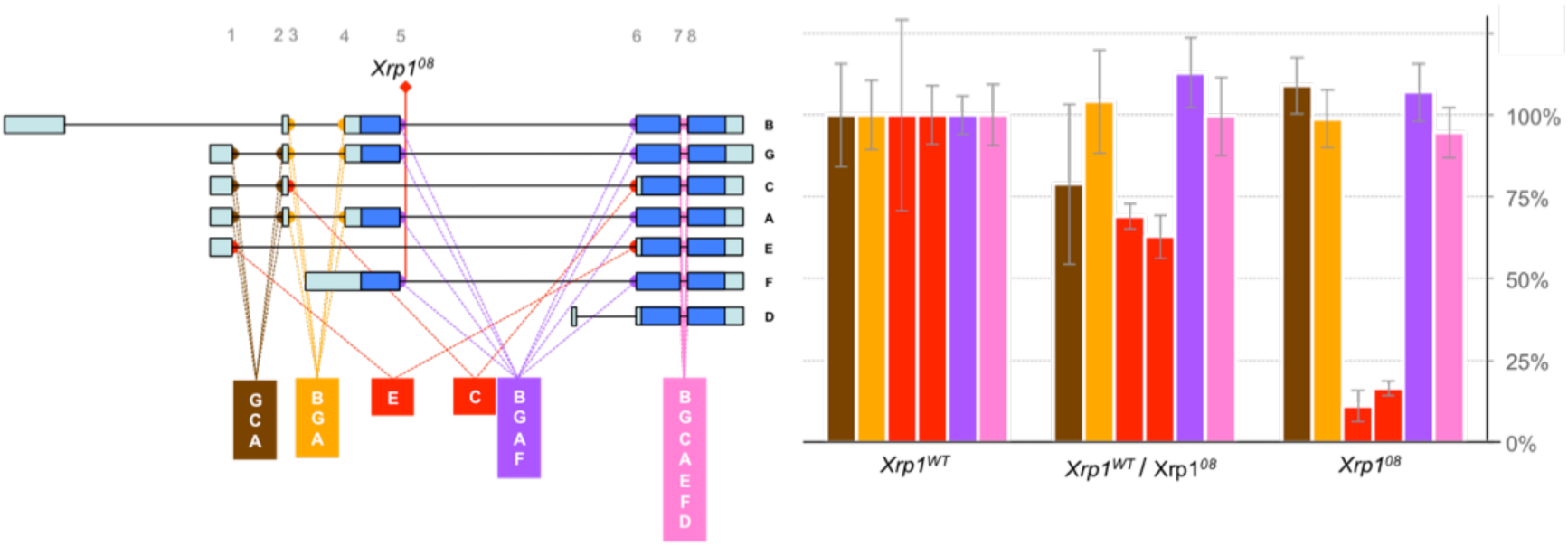
Intronic mutations reduce *Xrp1* transcripts E and C abundance. *Xrp1* transcripts are designated with a single letter from A to G on the left side. *Xrp1^08^* leads to a constitutive downregulation of the transcripts E and C, 9-folds and 6-folds respectively. (Left) Primers used for the qRT-PCR are numbered from 1 to 8. The exonic junctions amplified from each of these primer pairs is color-coded (1-2: brown, 3-4: orange, 1-6: red, 3-6: red, 5-6: purple and 7-8: magenta). The transcripts containing these exonic junctions are indicated in each colored box. (Right) qRT-PCR results are normalized to wild type. Error bars indicate standard deviations. Note that E and C transcript levels (red bars) diminish as the number of *Xrp1* mutant copies increases.

**Table S1.**
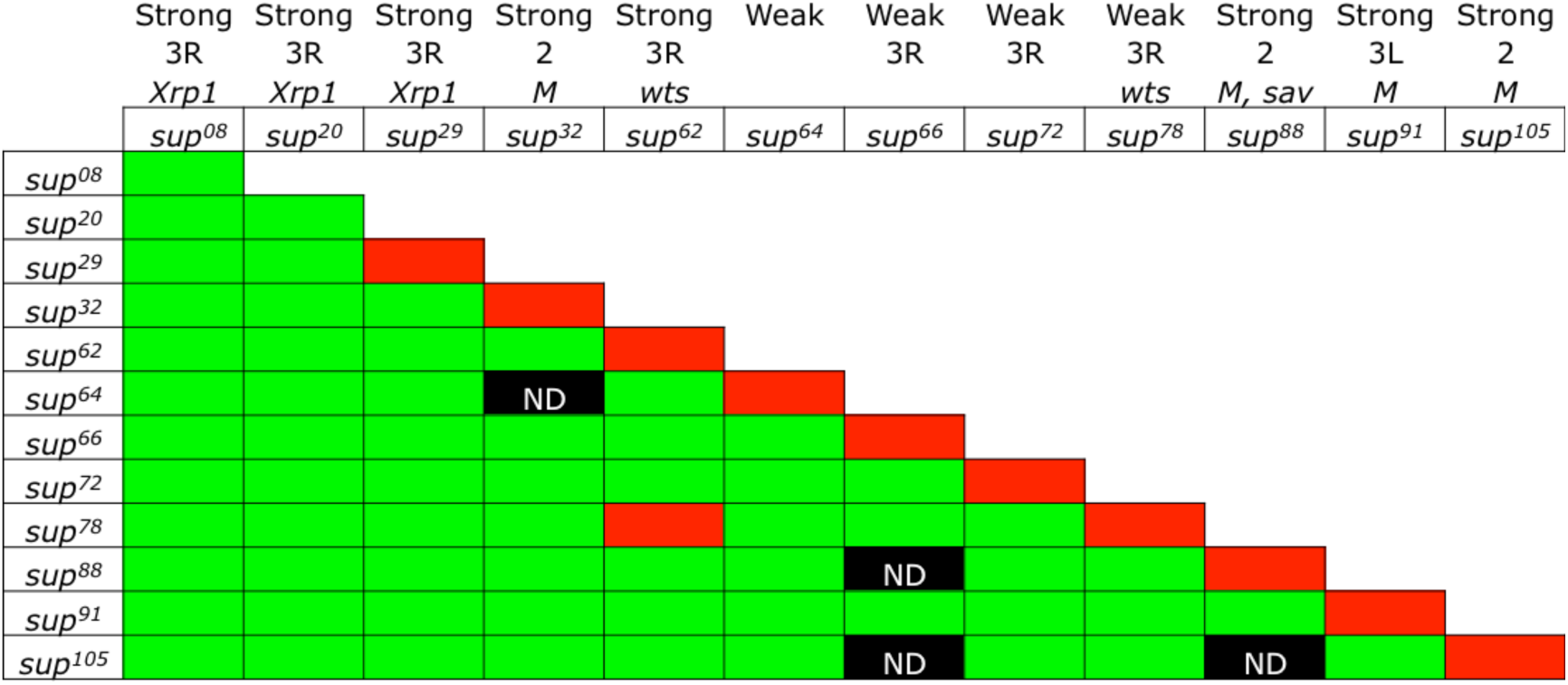
Complementation matrix of the EMS induced suppressors. Green color indicates complementation, red indicates non-complementation, ND stands for “not determined”. The upper line of the table indicates the strength of the suppression observed with the corresponding allele. The second line indicates the location of the suppressive mutation: “2” second chromosome, “3L” left arm of the third chromosome and “3R” right arm of the third chromosome. The third line indicates the mutated genes as far as they were identified, “M” indicating *Minute* phenotype. The fourth line indicates the names of the various suppressors.

**Figure S3.**
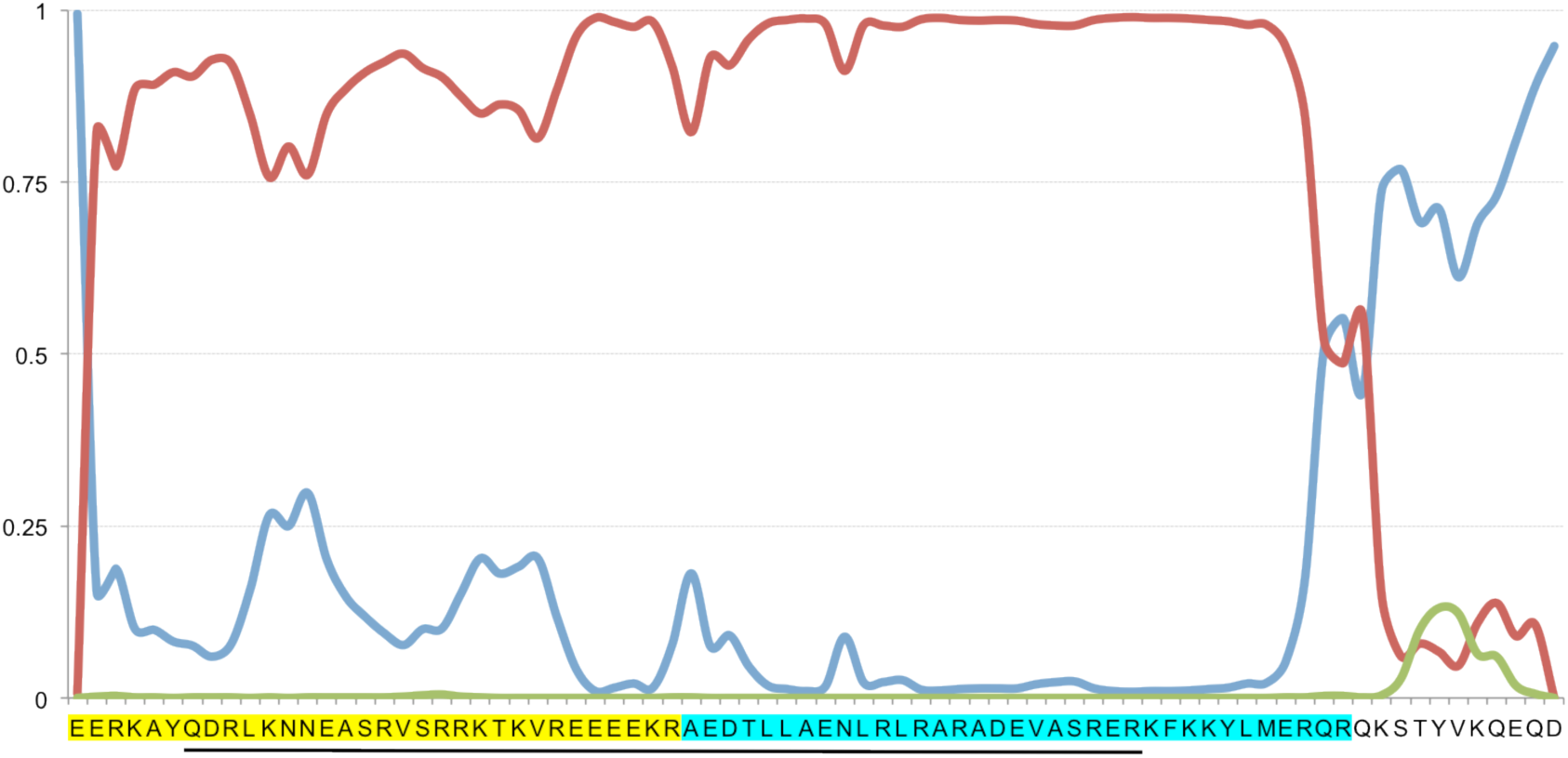
Structural conservation of the b-Zip domain of Xrp1. Secondary structure prediction (PSIPRED v3.3) of the C-terminal end of Xrp1. The conserved sequence of the bZip domain is predicted to adopt the expected alpha helix conformation (red line) necessary for the functionality of this domain. The blue line indicates the probability for a given AA to be in a random coil state and the green line indicates the probability for a given AA to be in a strand state. The black line delineates the sequence sharing a 40% identity with the human C/EBPs (PSI-BLAST)

**Figure S4.**
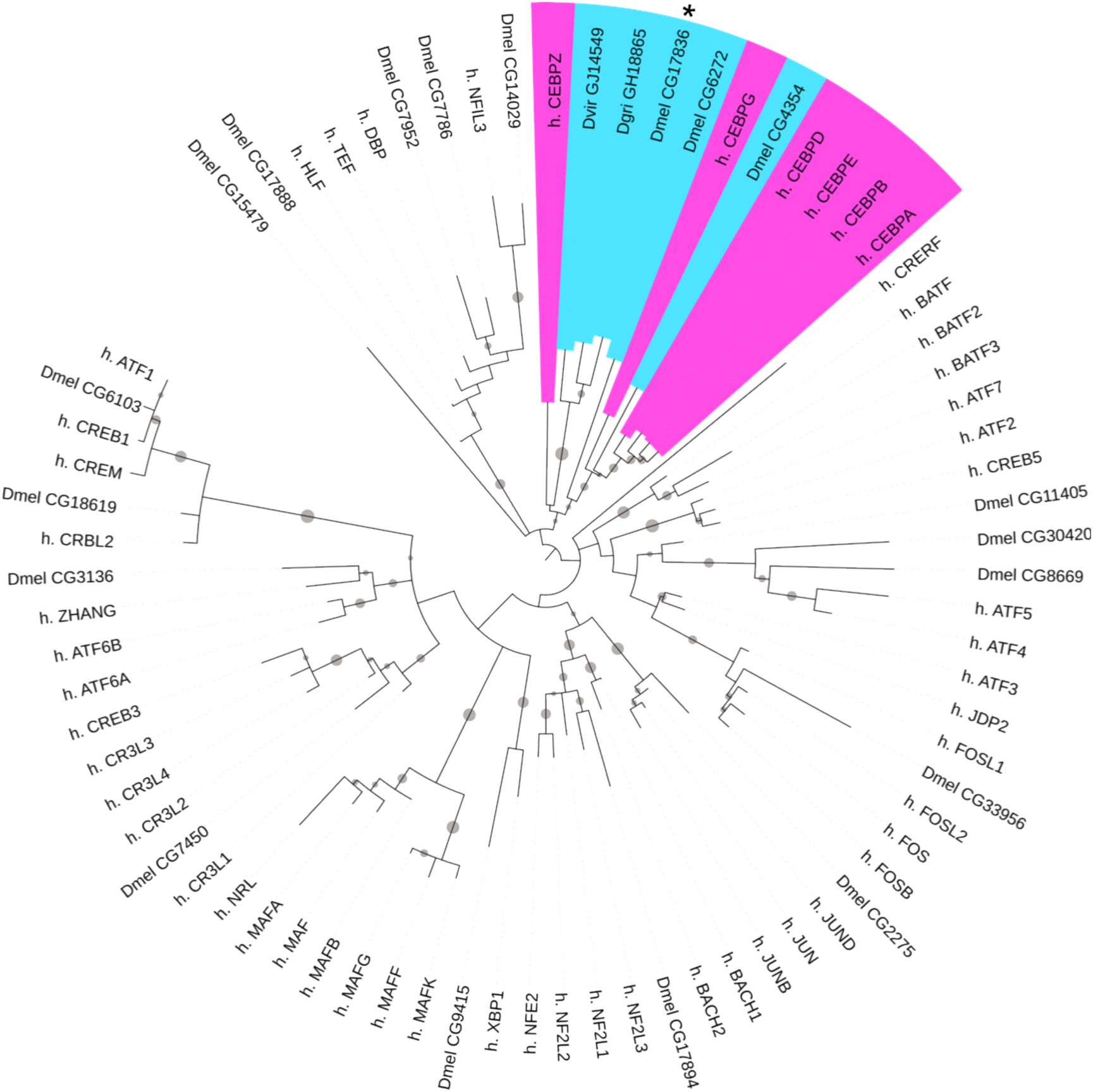
Xrp1 is homologous to the human C/EBPs. Phylogenetic reconstruction using the PhyML algorithm and visualized with the iTOL online tool branches with a bootstrap value superior to 80% are marked with a grey round sign. Humans C/EBPs are highlighted in magenta and *Drosophila*’s C/EBPs homologues are highlighted in blue. The asterisk marks the position of Xrp1.

**Table S2.** Protective function of Xrp1. Proportions are compared using the Z-test, significance level is set at 0.05. <u>Note</u>: At 60 μJ UVC only 7 Xrp1 homozygous mutants survived compared to 148 for the wild type control. This low number (i.e. 7) makes the statistical analysis on phenotypes at this relatively high dose irrelevant due to a higher sensitivity to sampling bias.

| | UVC<br>0 μJ | UVC<br>30 μJ | UVC<br>60 μJ |
| --- | --- | --- | --- |
| Number of yw L1 deposited on agar plates | 200 | 600 | 600 |
| Number of yw eclosed adults | 130 | 478 | 331 |
| Proportion of yw survival after UC exposure | 0.650 | 0.797 | 0.552 |
| Number of Xrp1(61) L1 deposited on agar plates | 200 | 600 | 600 |
| Number of Xrp1(61) eclosed adults | 127 | 230 | 7 |
| Proportion of Xrp1(61) survival after UC exposure | 0.635 | 0.383 | 0.012 |
| p value is inferior to | 0.757 | 1.0E-05 | 1.0E-05 |
| Statistical significance | No | Yes | Yes |
| Number of yw eclosed adults | 130 | 478 | 331 |
| Number of yw with an obvious adult phenotype | 17 | 72 | 148 |
| Proportion of yw with an obvious adult phenotype | 0.131 | 0.151 | 0.447 |
| Number of Xrp1(61) eclosed adults | 127 | 230 | 7 |
| Number of Xrp1(61) with an obvious adult phenotype | 20 | 96 | 4 |
| Proportion of Xrp1(61) with an obvious adult phenotype | 0.157 | 0.417 | 0.571 |
| p value is inferior to | 0.542 | 1.0E-05 | 0.513 |
| Statistical significance | No | Yes | No |

**Figure S5.**
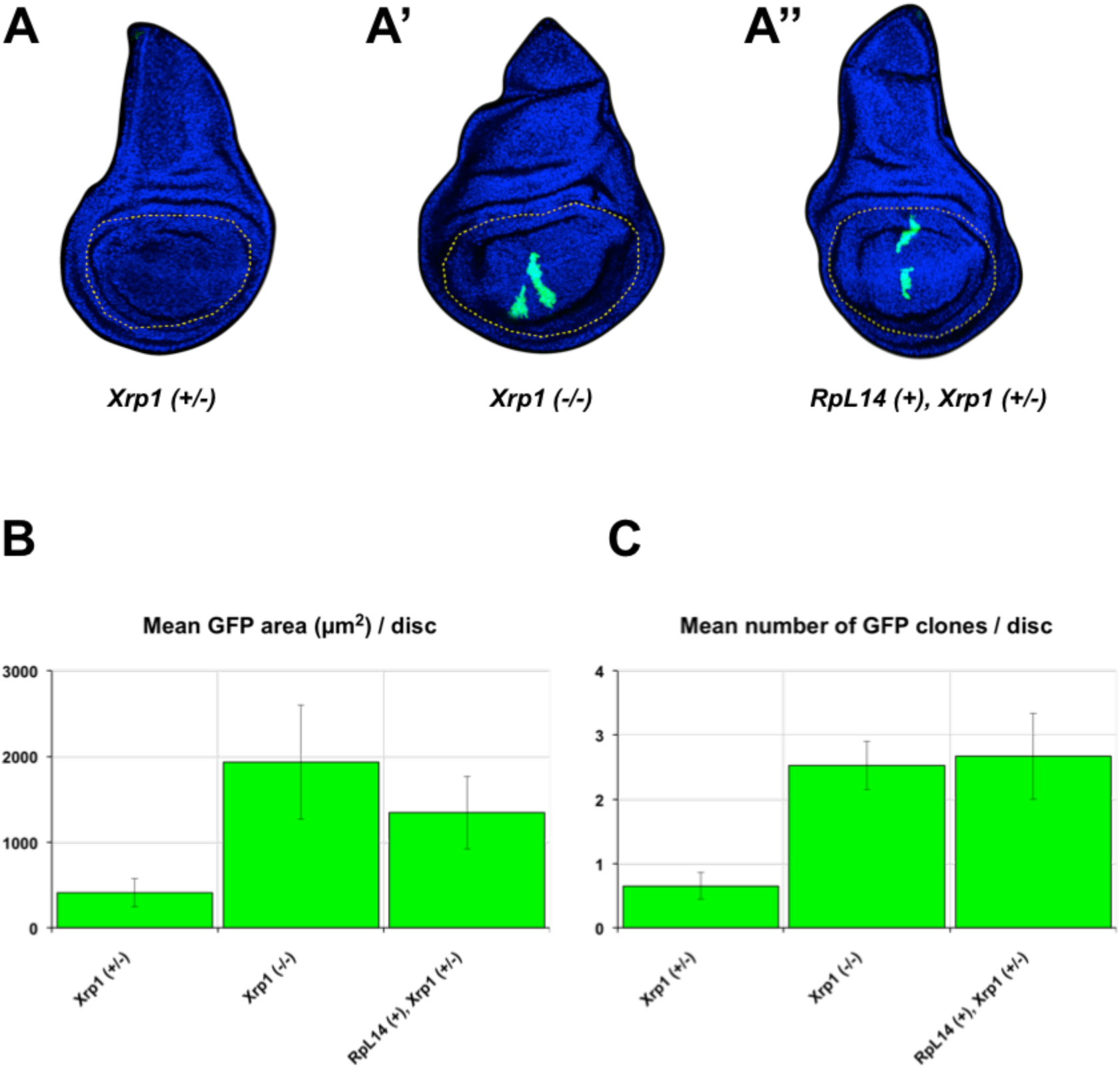
Loss of *Xrp1* function suppresses RpL14^+/−^ loser cell eliminations. Removal of one copy of the *RpL14* gene generates *RpL14* heterozygous loser clones that are efficiently eliminated (Meyer et al., 2014). Induction of mitotic recombination excises a copy of the *RpL14* gene and places the expression of GAL4 under the control of the wing pouch specific *Salm* enhancer, resulting in GFP expressing *RpL14*^+/−^ loser clones in the wing pouch region (Meyer et al., 2014). The wing pouch region is outlined with a dashed orange line. **(A)** Loser clones heterozygous mutant for *Xrp1* are efficiently eliminated (n= 20). **(A’)** In contrast, loser clones homozygous mutant for *Xrp1* are not eliminated (n=23). **(A’’)** Similarly, loser clones (heterozygous mutant for *Xrp1*) in which the *RpL14* deficit is restored via *RpL14* overexpression are not eliminated (n=9). **(B)** Quantification of the total GFP area (μm^2^) per disc, and **(C)** quantification of the number of clones per disc reveals a suppressive potential for *Xrp1* deficient clones that is similar to that of *RpL14* supplemented clones (Mann-Whitney test). Bars on the plot represent SEM. Detailed genotypes for each figure panel are listed in Supplementary Table S3.

**Figure S6.**
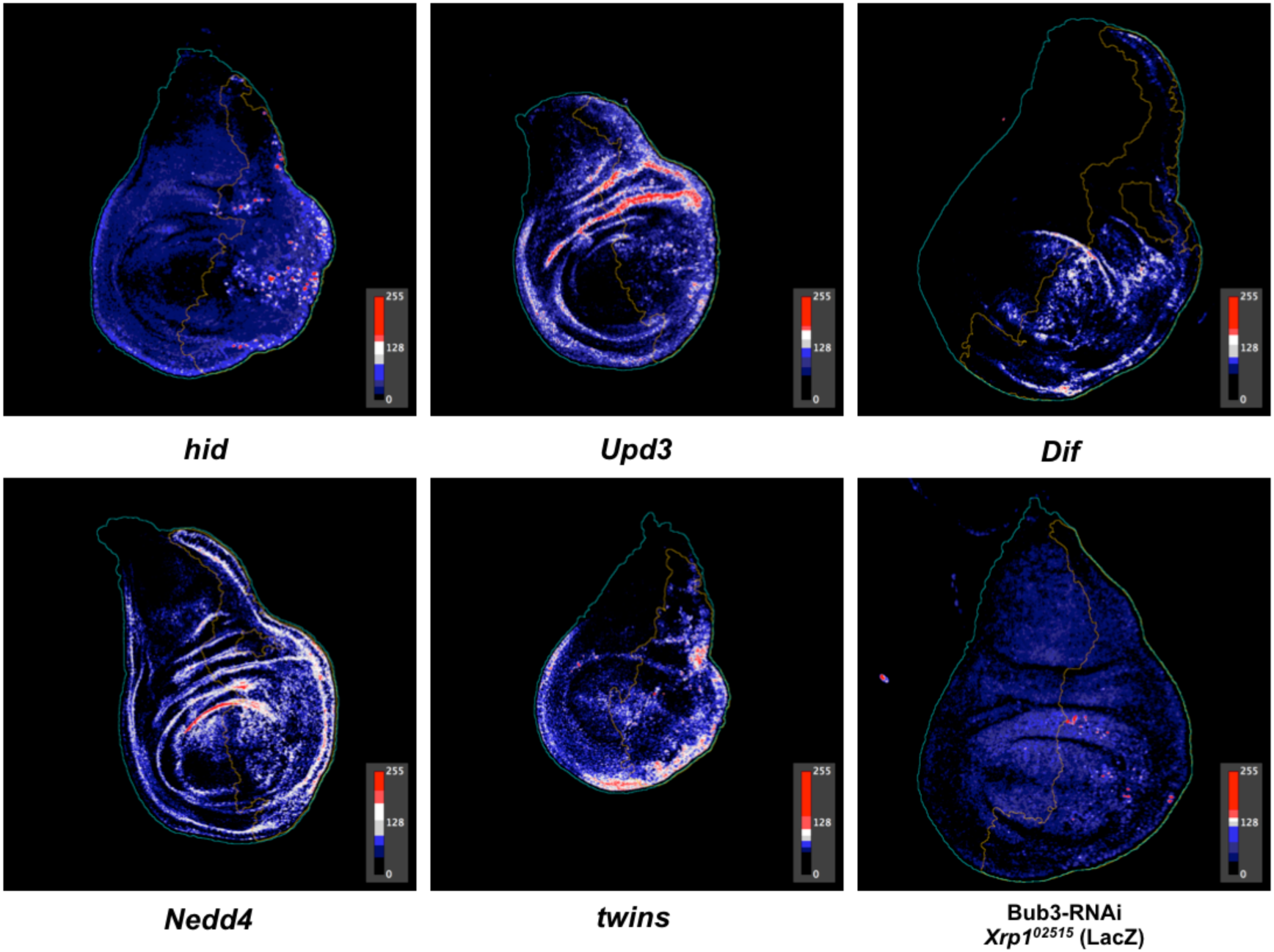
LUT processed images from Fig. 3E and Fig. 4C. LUT (lookup table) processed images where gray scale intensity values are mapped to a desired color space (blue is minimu, red is maximum).

**Figure S7.**
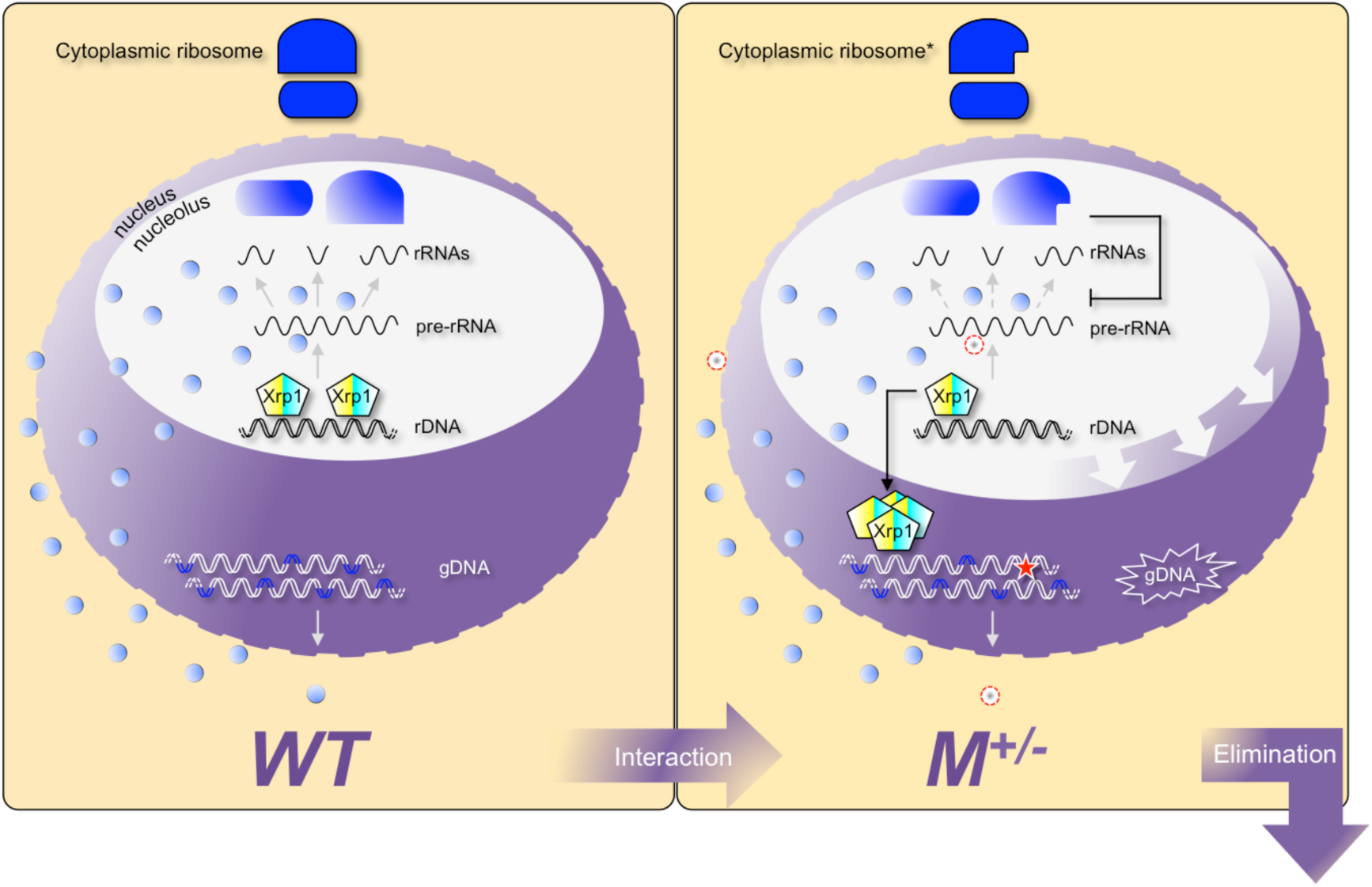
Working hypothesis on the possible role of the RPs-Xrp1 axis in the maintenance of genomic integrity. In wild-type (WT) cells, there is a complete set of ribosomal protein genes (*RPGs*) (regions colored in blue on the genomic DNA). Under normal conditions, Xrp1 binds to the ribosomal DNA (rDNA) and would therefore be sequestered in the nucleolus (white ellipse). Loss of genomic integrity in these cells can be revealed by the loss of at least one *RPG* (red star). In these *M*+/- cells one of the ribosomal proteins becomes limiting (dashed red circles indicate absence) which stalls ribosome production (barheaded black line) in the nucleolus. This results in the accumulation of unprocessed rRNAs (pre-RNA) that enlarge the nucleolus (large white arrows). Xrp1 is released (arrow-headed black line) into the nucleus (purple circle) where it amplifies and initiates its transcriptional program leading to cell elimination.

**Figure S8.**
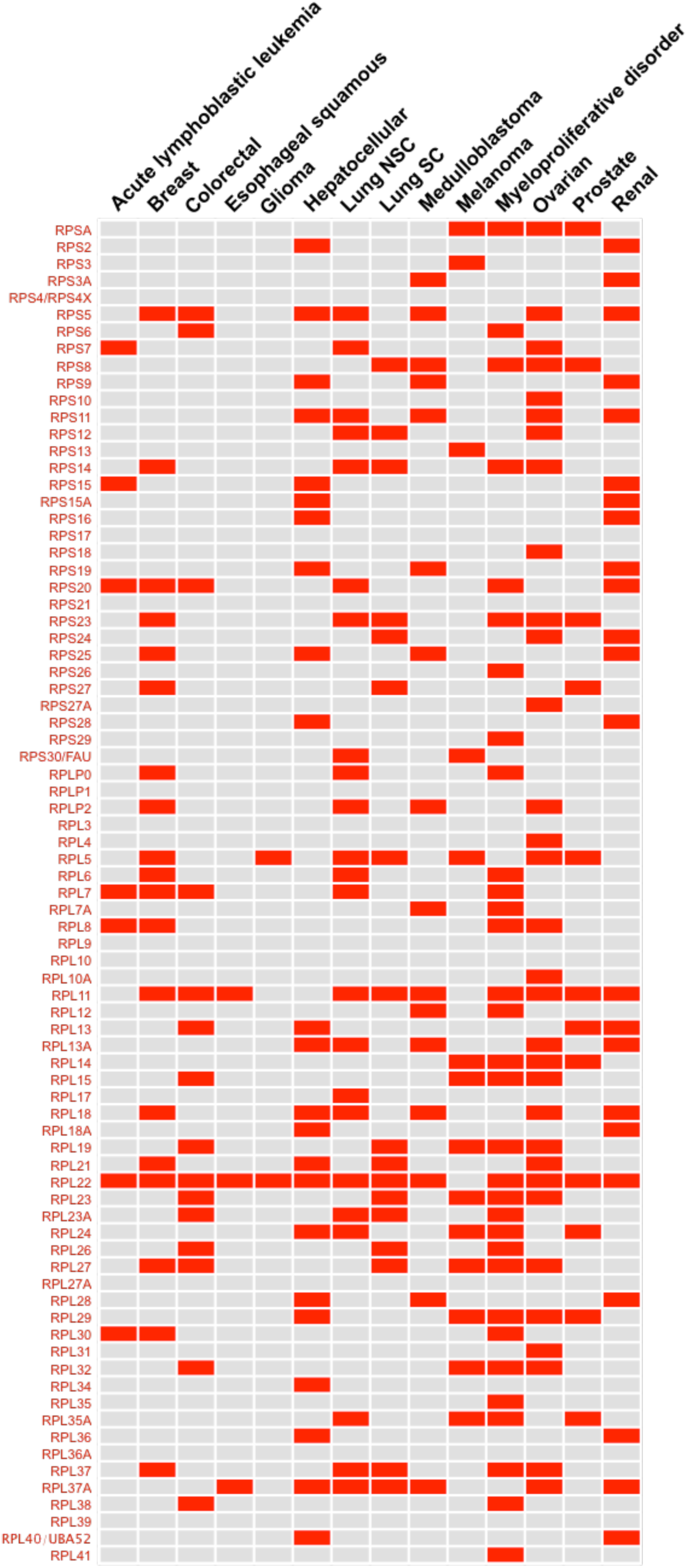
*RPG* deletion profile in human cancer. Human ribosomal protein genes are also distributed across the genome. As a result, these genes are often deleted due to genomic instability. Lanes correspond to each of the 79 human RPGs and columns correspond to specific types of cancer. Red indicates that a particular RPG is significantly deleted according to the genome-wide GISTIC analysis results publically available at the Tumorscape portal (Beroukhim et al., 2011).

**Table S3:**
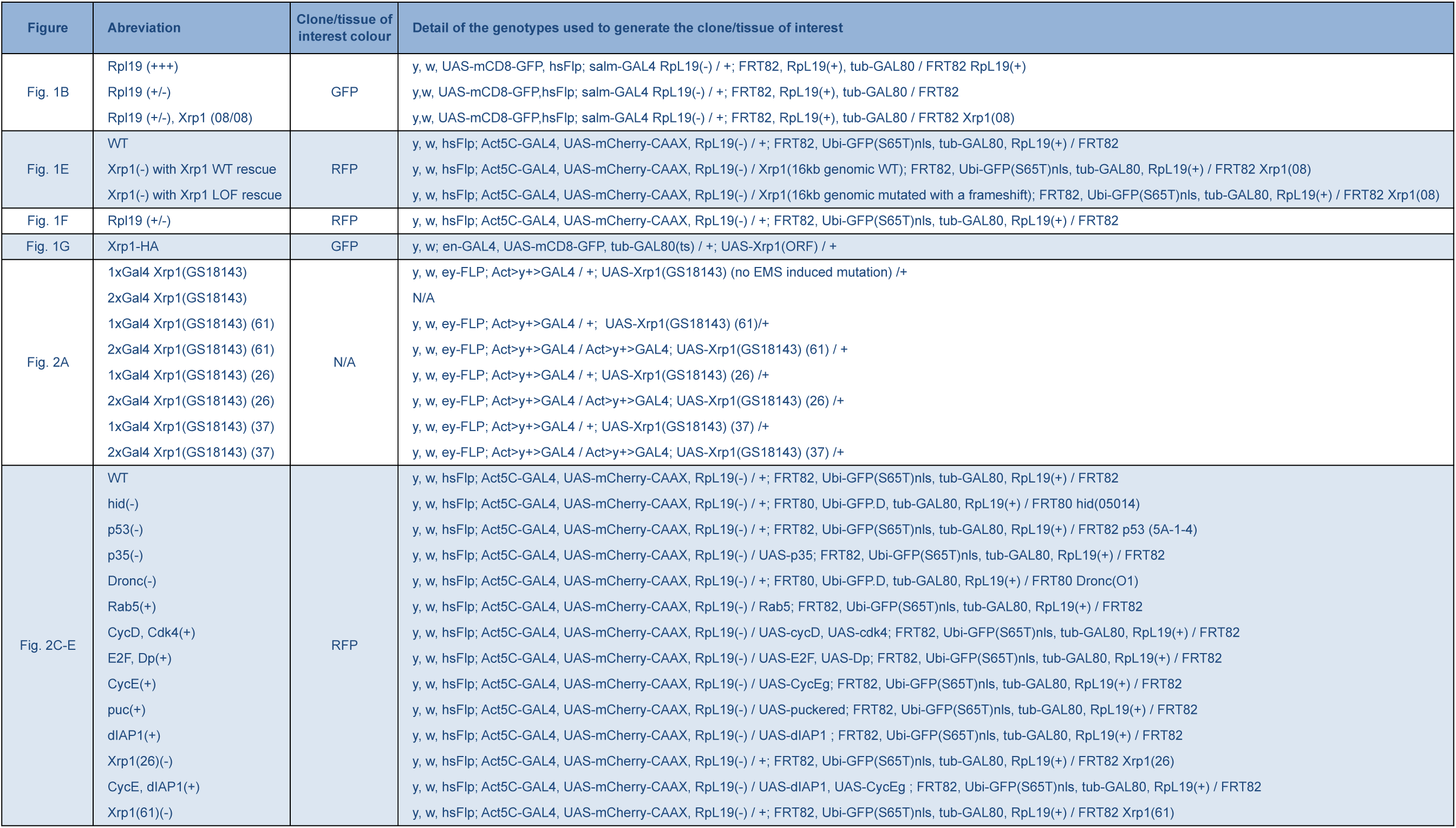
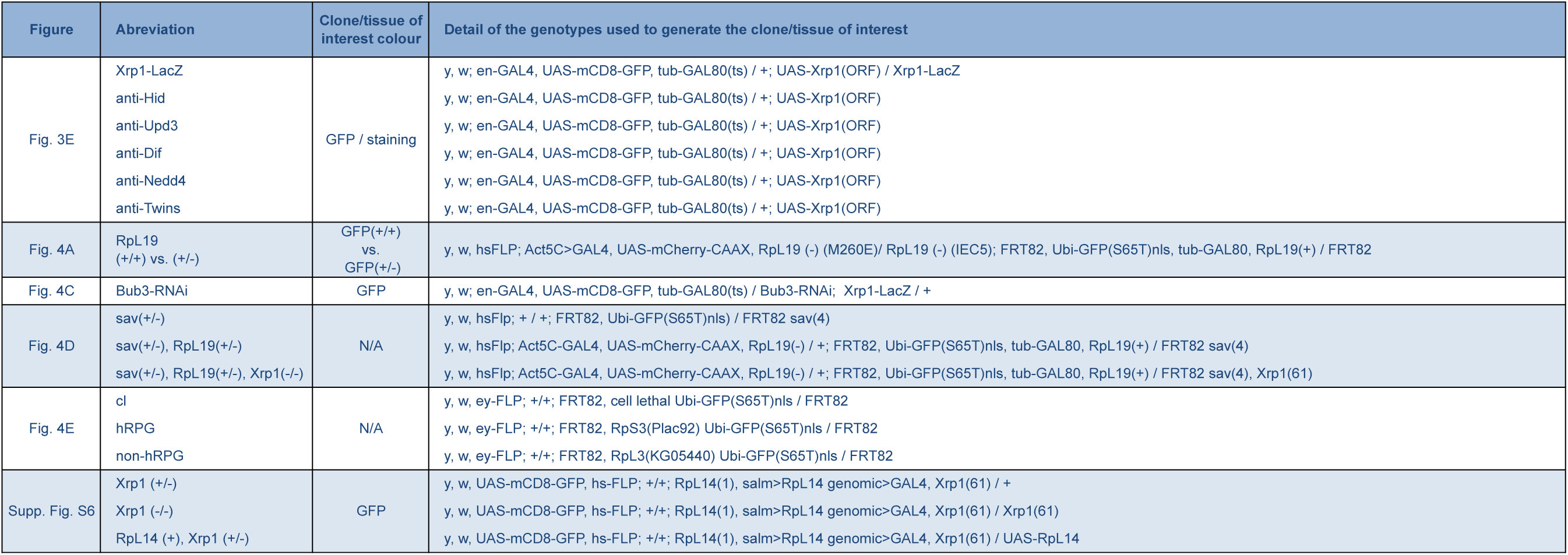

